# Brain injuries and complex motor learning suppress Olig2 in a subpopulation of oligodendrocyte precursor cells

**DOI:** 10.1101/2022.06.18.496631

**Authors:** Li-Pao Fang, Qing Liu, Erika Meyer, Anna Welle, Wenhui Huang, Anja Scheller, Frank Kirchhoff, Xianshu Bai

**Author notes:** To whom correspondence should be addressed: Frank Kirchhoff: Molecular Physiology, Center for Integrative Physiology and Molecular Medicine, Building 48, University of Saarland, 66421 Homburg, Germany ORCID: 0000-0002-2324-2761 Xianshu Bai: Molecular Physiology, Center for Integrative Physiology and Molecular Medicine, Building 48, University of Saarland, 66421 Homburg, Germany ORCID: 0000-0002-4758-1645. **Li-Pao Fang:**, **Qing Liu:**, **Erika Meyer:**, **Anna Welle:**, **Wenhui Huang:**, **Anja Scheller:**. **Author contributions:** LF and XB conceived and designed the experiments; LF induced SWI model; XB performed kainate injection; EM performed MCAO surgery; LF, QL and WH performed Erasmus Ladder experiments. LF and QL carried out slice preparation, immunohistochemistry, confocal imaging and data analysis. AS performed AxioScan imaging. AW performed analysis of single cell RNA transcriptomic data. XB and FK supervised the project; XB wrote the manuscript with input from the other authors. **Data availability** The raw data are available from the corresponding authors upon reasonable request. Data of single cell RNA sequencing was submitted with the manuscript in Supplementary file. **Abbreviations:** bHLH: basic helix-loop-helix; Cspg4: chondroitin sulfate proteoglycan 4; dpi: days post injury; CCA: common carotid artery; MCAO: middle cerebral artery occlusion; LPS: lipopolysaccharide; Olig2: oligodendrocyte transcription factor 2; OPCs: oligodendrocyte precursor cells; p: postnatal day; PDGFRα: platelet derived growth factor receptor alpha; SWI: stab wound injury.

## Abstract

Oligodendrocyte precursor cells (OPCs) are uniformly distributed in the mammalian brain, however their function is rather heterogeneous in respect to their origin, location, receptor/channel expression and age. The basic helix-loop-helix transcription factor Olig2 is expressed in all OPCs as a pivotal determinant of their differentiation. Here, we identified a subset (2-26%) of OPCs lacking Olig2 in various brain regions including cortex, corpus callosum, CA1 and dentate gyrus. These Olig2 negative (Olig2^neg^) OPCs were enriched in the juvenile brain and decreased subsequently with age, being rarely detectable in the adult brain. However, the loss of this population was not due to apoptosis or microglia-dependent phagocytosis. Unlike Olig2^pos^ OPCs, these subset cells could not be labelled for the mitotic marker Ki67. And, accordingly, BrdU was incorporated only by a three-day long-term labeling but not by a two-hour short pulse, suggesting these cells do not proliferate any more but were derived from proliferating OPCs. The Olig2^neg^ OPCs exhibited a less complex morphology than Olig2^pos^ ones. Olig2^neg^ OPCs preferentially remain in a precursor stage rather than differentiating into highly branched oligodendrocytes. Changing the adjacent brain environment, e.g. by acute injuries or by complex motor learning tasks stimulated the transition of Olig2^pos^ OPCs to Olig2^neg^ cells in the adult. Taken together, our results demonstrate that OPCs transiently suppress Olig2 upon changes of the brain activity.

**Table of Contents Image:** 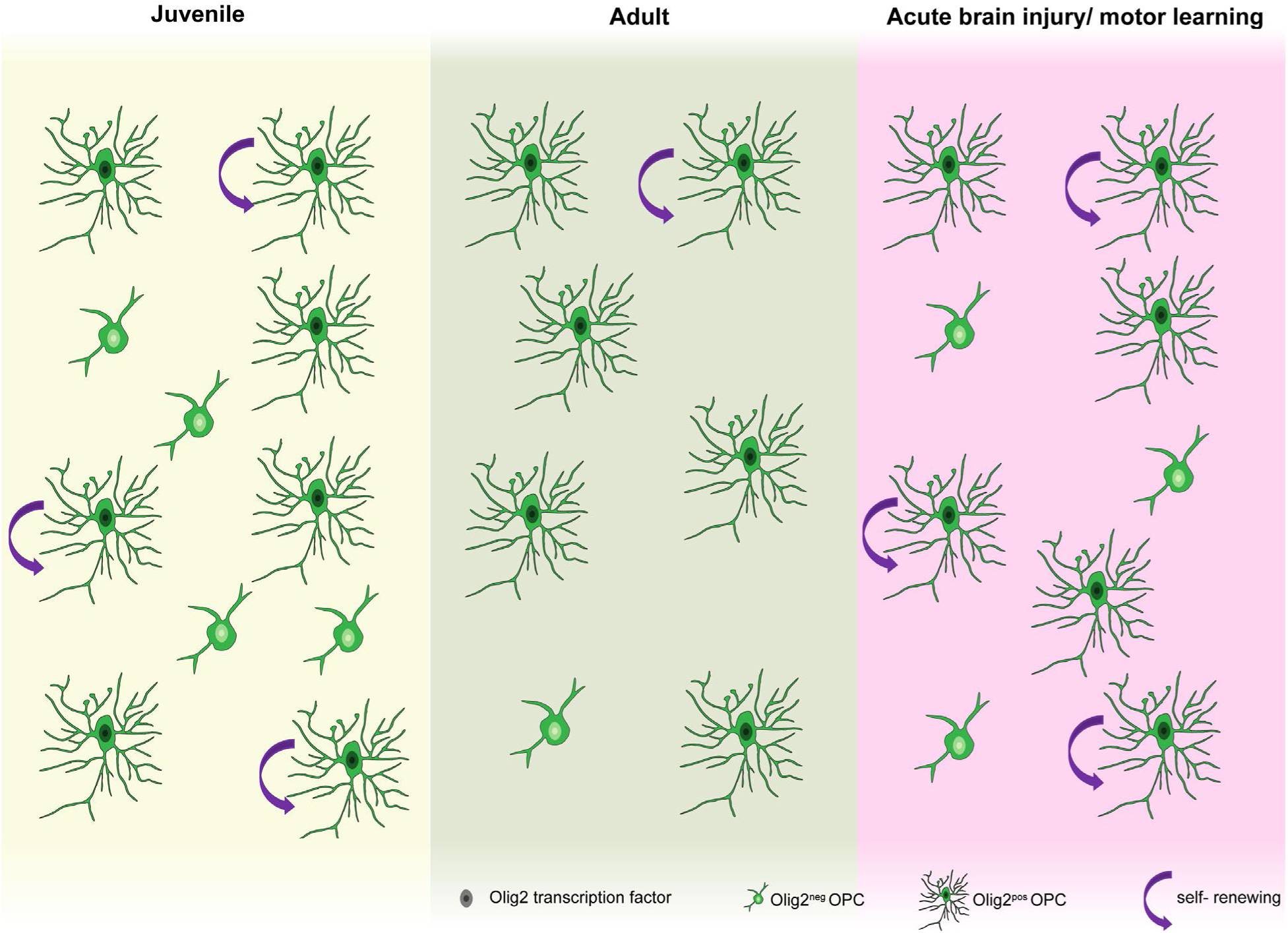

**Main points:** A subset of OPCs do not express Olig2, of which population peaks in the juvenile brain while wanes with age. Plastic changes of the brain by acute injuries or complex motor learning stop the expression of Olig2 in OPCs.

## Introduction

Oligodendrocyte precursor cells (OPCs) are widely distributed within the central nervous system (CNS), however they are rather a heterogeneous population regarding to their location, origin, age and function (Marisca et al., 2020; Marques et al., 2018; Marques et al., 2016; Spitzer et al., 2019; Viganò, Möbius, Götz, & Dimou, 2013). A conserved feature of OPCs is the capability to give rise to oligodendrocytes throughout the life (Trotter, Karram, & Nishiyama, 2010). OPC differentiation is tightly regulated by intrinsic and extrinsic factors (Rowitch & Kriegstein, 2010). The basic helix-loop-helix transcription factor Olig2, is one of the pivotal intrinsic determinants for oligodendrocyte specification (Liu et al., 2007; Q. R. Lu et al., 2002; Q. R. Lu et al., 2000; Maire, Wegener, Kerninon, & Nait Oumesmar, 2010; Zhou & Anderson, 2002; Zhou, Wang, & Anderson, 2000). In Olig2 null mice, OPC formation fails at embryonic and perinatal stages (Ligon et al., 2006; Mei et al., 2013), while overexpression of Olig2 triggers OPC differentiation and precocious myelination (Maire et al., 2010; Wegener et al., 2015). Although Olig2 is widely expressed throughout the oligodendroglial development, its function is rather specific to the cell stage (Mei et al., 2013). For instance, Olig2 expression in OPCs promotes cell differentiation and subsequent myelination, while in newly formed oligodendrocytes Olig2 seems to suppress maturation and myelination. Apart from lineage commitment, Olig2 exerts function in OPC migration. Overexpression of Olig2 accelerates OPC migration, differentiation and subsequently promoted remyelination in the lysolecithin model of multiple sclerosis (Wegener et al., 2015).

Despite of the importance and abundance of Olig2 for the lineage of oligodendrocytes, a proportion of NG2^pos^ cells could not be detected for Olig2 expression in the healthy perinatal (about 1 %) and adult cortex (8-30 %) (Buffo et al., 2005; Ligon et al., 2006). This NG2^pos^Olig2^neg^ population was even larger after a stab wound injury (Buffo et al., 2005). These NG2^pos^Olig2^neg^ cells were defined according to their immunoreactivity to NG2 antibody. Of note, NG2 is also expressed by pericytes under physiological conditions and by a small population of microglia triggered by acute brain injuries (W. Huang, Bai, Meyer, & Scheller, 2020; W. Huang et al., 2014). In addition, due to the cleavage of the extracellular domain of NG2 protein under pathological conditions, NG2 immunoreactivity per se cannot be referred as NG2 expression. Therefore, it is yet elusive whether or not Olig2^neg^ OPCs exist in the brain. If yes, how does the population develop with age? Are they functionally different to the Olig2^pos^ OPCs?

To address these open questions, we distinguished OPCs from other NG2 expressing cells with immunostaining of platelet derived growth factor receptor α (PDGFRα), a well-established marker of OPCs (Nishiyama, Lin, Giese, Heldin, & Stallcup, 1996). We analyzed different brain regions, e.g. cortex, corpus callosum, CA1 and dentate gyrus, at different ages ranging from postnatal day (p) 5 to 44 weeks. PDGFRα^pos^Olig2^neg^ population was progressively increased at the first two postnatal weeks, followed by a consistent decline. Derived from Olig2^pos^ cells, Olig2^neg^ cells were never expressing Ki67 and exhibited a less complex morphology. Complex motor learning or acute brain injuries, which are known to stimulate brain activity change, triggered increased the population of Olig2^neg^ OPCs in the adult hippocampus and cortex, respectively. In summary, our study demonstrates that OPCs suppress Olig2 expression upon the change of brain activity.

## Results

### A subset of OPCs does not express Olig2

To investigate whether Olig2-non-expressing OPCs exist or not, we performed immunostaining of PDGFRα (Pα) and Olig2 in coronal section of healthy mouse brain (Fig. 1A). At postnatal day (p) 14, we observed a small population of Pα^pos^ cells expressing low level or no Olig2 in various brain regions, including primary motor cortex (MOp, Fig. 1B-D), corpus callosum (CC, Fig. 1E), CA1 (Fig. 1F) and in the dentate gyrus region (DG, Fig. 1G). To further characterize these cell subtypes, we classified Pα^pos^ cells into Olig2^pos^ cells with high immunoreactivity (fluorescence intensity (FI)>10^3^, 96.52 %) and Olig2^neg^ cells with no Olig2 expression (FI<10^3^, 3.48 %) (Supplementary Fig. 1A). These results indicate that indeed a subset of OPCs do not express Olig2 in several brain regions.

**Figure 1.**
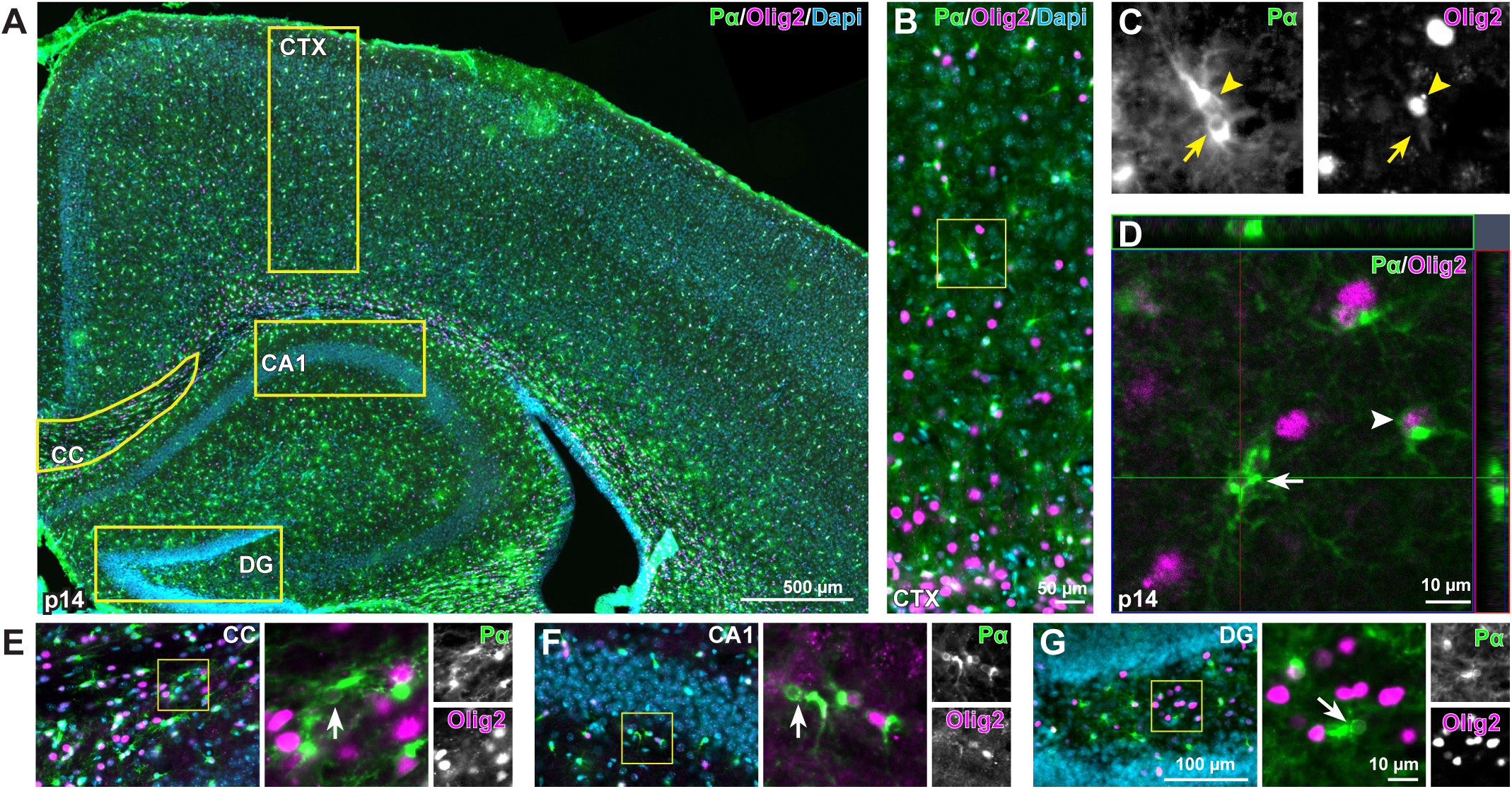
A subset of PDGFRα-positive cells do not express Olig2. **A**, Overview of PDGFRα (Pα, green) and Olig2 (magenta) immunostaining of coronal brain slices from p14 mice. **B, E-G,** Representative images of cortex (CTX), corpus callosum (CC), CA1 and dentate gyrus (DG) stained with Pα and Olig2. **C,** Magnified views of Pα^pos^Olig2^neg^ cells from **B** (boxed area). **D,** Confocal images showing Pα^pos^ cells without Olig2 expression in cortex. Arrowheads: Olig2^pos^ OPCs, arrows: Olig2^neg^ OPCs.

To confirm these Pα^pos^Olig2^neg^ cells are OPCs, we further performed immunostainings by combining Pα and Olig2 with various cell type specific markers at p14, when the population peaks. All Pα^pos^Olig2^neg^ cells were immunopositive for NG2 with bona fide OPC morphology (Fig. 2A, Supplementary Fig. 2A). However, the majority (about 98 %) of Pα^pos^Olig2^neg^ cells were also negative for Sox10 (transcription factor, another lineage marker of oligodendrocytes) (Fig. 2B, Supplementary Fig. 2B). In addition, these cells never expressed markers of mature oligodendrocytes (e.g. APC CC1, Fig. 2C, Supplementary Fig. 2C), astrocytes (GFAP, Fig. 2D, Supplementary Fig. 2D; GS, Fig. 2E, Supplementary Fig. 2E), microglia (Iba1, Fig. 2F, Supplementary Fig. 2F), neurons (NeuN, Fig. 2G, Supplementary Fig. 2G), or precursors (Sox2, Fig. 2H; Supplementary Fig. 2H). Therefore, our results suggest that Pα^pos^Olig2^neg^ cells are OPCs/NG2 glia.

**Figure 2.**
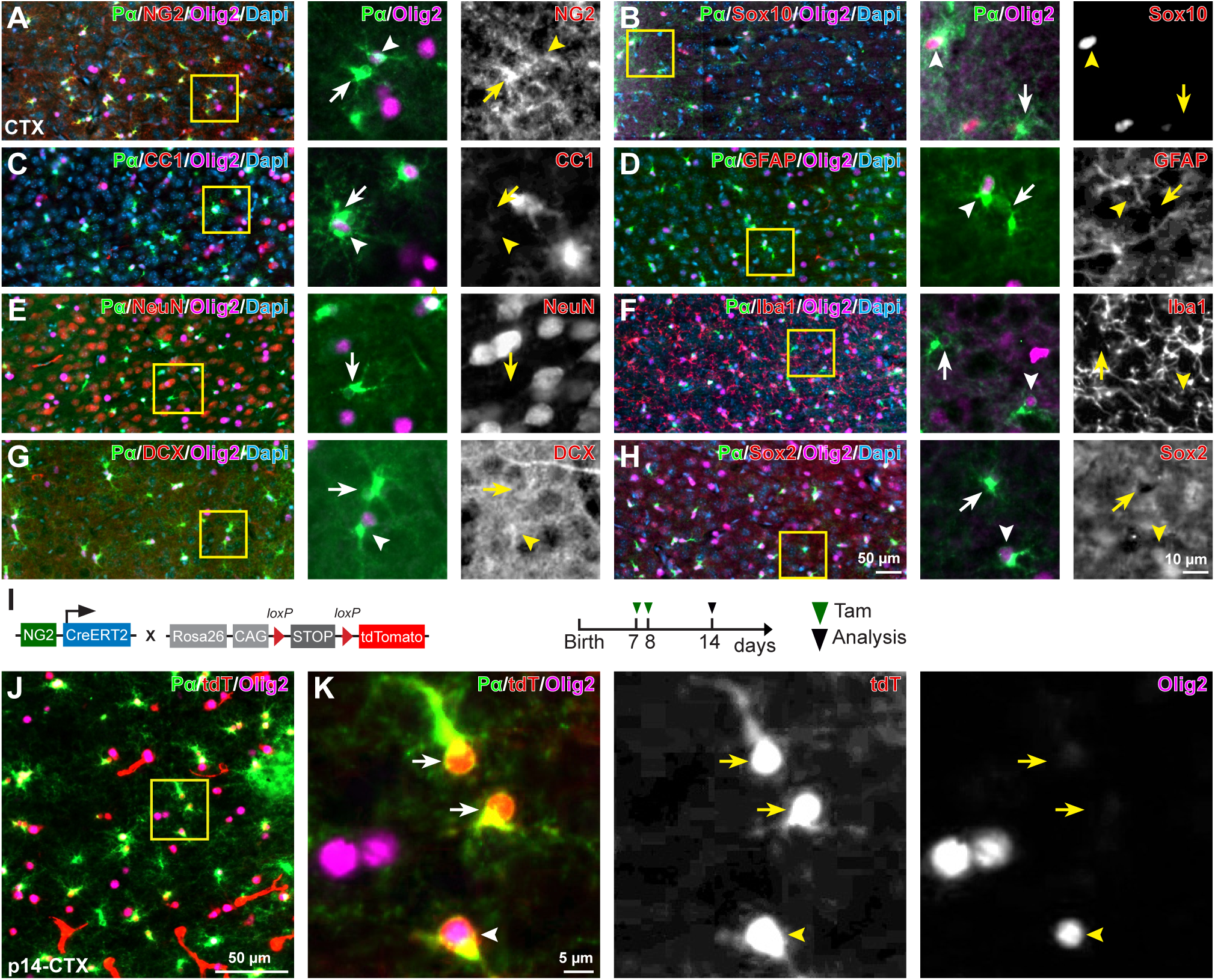
PDGFRα^pos^Olig2^neg^ cells are *bona fide* OPCs. **A, B,** Pα^pos^Olig2^neg^ cells are immunopositive for NG2 (**A**), but rarely express Sox10 (**B**). **C-H**, Pα^pos^Olig2^neg^ cells are not oligodendrocytes (**C**), astrocytes (**D, E**), microglia (**F**), neurons (**G**) or neural precursors (**H**). **I**, scheme of transgene construct and experimental schedule. **J**, Immunostaining of Pα and Olig2 in the cortex of NG2-CreER^T2^ x R26-tdTomato mice at p14. **K**, Magnified views of Pα^pos^Olig2^neg^ cells expressing tdTomato (tdT) from the area indicated in **J**. Arrowheads: Olig2^pos^ OPCs, arrows: Olig2^neg^ OPCs.

To further substantiate the lack of Olig2 in a subset of OPCs, we took advantage of NG2-CreER^T2^ x R26-lsl-tdTomato mice, in which OPCs, their descendent oligodendrocytes as well as pericytes express tdTomato (tdT) after tamoxifen induced recombination. Tamoxifen was administrated at p7 and 8 (Fig. 2I), inducing tdT expression in about 80 % of OPCs at p14 (Fang et al., 2022). Again, we observed a small population (about 3 %) of Pα^pos^tdT^pos^ cells expressing no Olig2 (Fig. 2J, K), confirming that a subset of OPCs do not express Olig2.

### Olig2^neg^ OPCs are enriched in the juvenile brain

To elucidate whether this cell population exists temporarily or over all ages under physiological conditions, we further analyzed the mice at p5, p9, p14/2 week (w), 4 w, 9 w, 11 w, 22 w and 44 w of age. In MOp, we observed about 1.3 ± 0.2 Olig2^neg^ cells/1x10^-3^ mm^3^ at p5 (Fig. 3A, G), i.e. about 2 % of all OPCs (Fig. 3H). Progressively, this Pα^pos^Olig2^neg^ cell population increased reaching 3.1 ± 0.8 cells/1x10^-3^ mm^3^ (accounting for 7 % of all OPCs) at p14 (Fig. 3G, H). The same increase was observed in other brain regions (Fig. 3G, Supplementary Fig. 1C-E) where Pα^pos^Olig2^neg^ cells were covering 9 %, 11 %, 13 % and 26 % of all OPCs in CC, mPFC, DG and CA1, respectively (Fig. 3H). After p14, the density of Pα^pos^Olig2^neg^ cells continuously decreased. It became rarely detectable in the adult cortex (MOp and mPFC), but it still remained at a rather high level in CC (Fig. 3D-F, G, H). Please note, the portion of Pα^pos^Olig2^neg^ cells among all Pα^pos^ cells increased till the fourth postnatal week (Fig. 3H), likely due to the reduced pool of Pα^pos^Olig2^pos^ cells during development as observed by us and others (Fig. 3I) (Kessaris et al., 2006). Overall, our results indicate that Olig2^neg^ OPCs are present in the brain throughout life, with higher population during the development.

**Figure 3.**
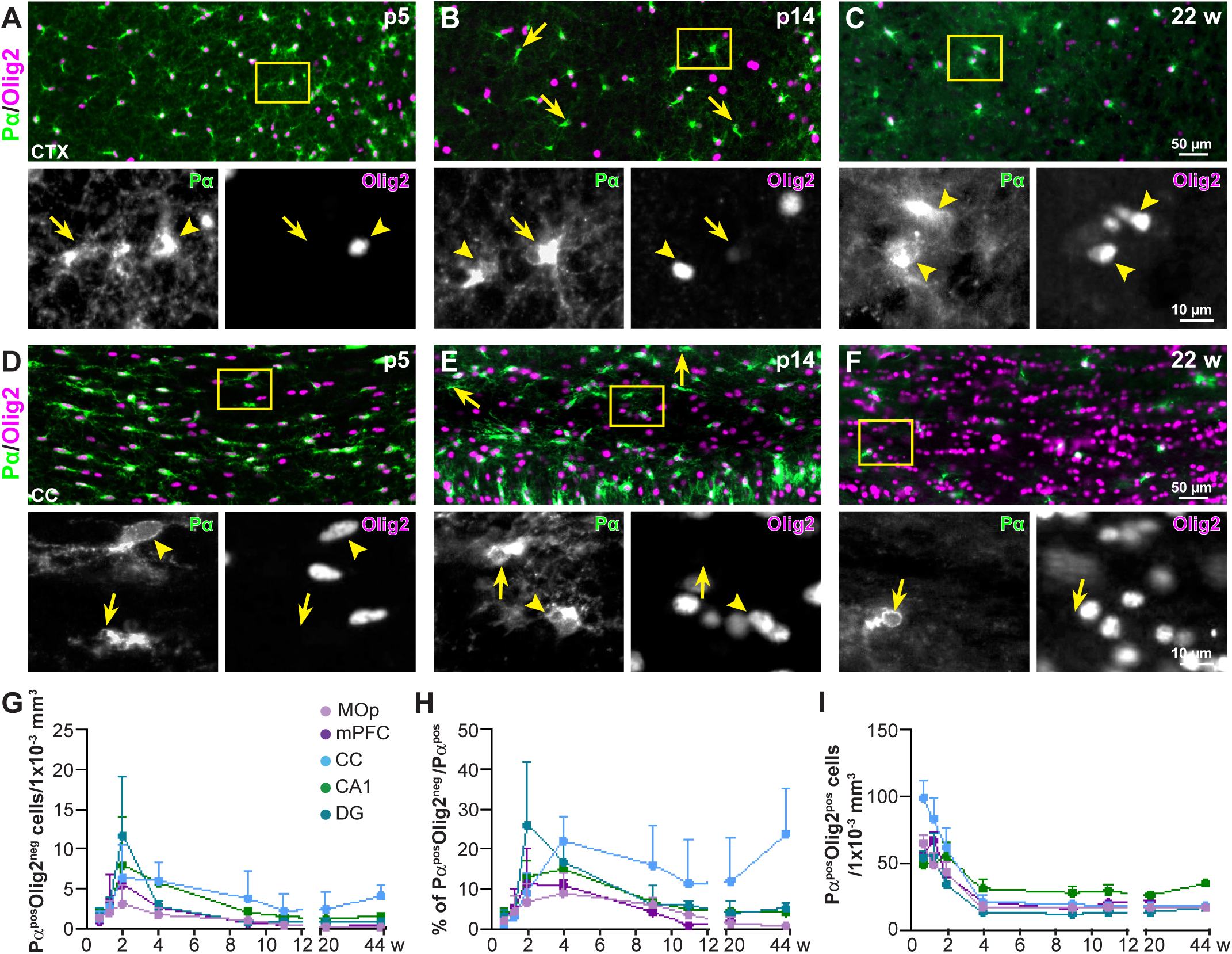
Olig2^neg^ OPCs are enriched in the juvenile brain. **A-C**, Immunostaining of Pα and Olig2 in the cortex of mice at the age of p5 (**A**), p14 (**B**) and 22 weeks (**C**). **D-F**, Immunostaining of Pα and Olig2 in the corpus callosum of mice at the age of p5 (**D**), p14 (**E**) and 22 weeks (**F**). **G-I,** Quantification of density and proportion of Pα^pos^Olig2^neg^ (**G, I**) and Pα^pos^Olig2^pos^ cells (**H**) reveals maximal population of Pα^pos^Olig2^neg^ cells in the juvenile brain (primary motor cortex, MOp; medial prefrontal cortex, mPFC; Corpus callosum, CC, CA1 and DG). Arrowheads: Olig2^pos^ OPCs, arrows: Olig2^neg^ OPCs.

### Olig2^neg^ OPCs exhibit low proliferative activity

To understand whether the dynamic of the population size was attributed to cell proliferation, cell death or to a transient regulation of Olig2 gene expression, we firstly analyzed apoptosis and phagocytosis of these cells by performing immunostaining of cleaved caspase 3 (CC-3, a well-established marker for apoptosis) (Fig. 4A) and CD68 (phagocytic marker, Fig. 4B) at p14. Notably, Olig2^neg^ OPCs were neither positive for CC-3 nor for CD68 (Fig. 4A, B, Supplementary Fig. 3), largely excluding apoptotic loss or microglia mediated elimination of Olig2^neg^ OPCs.

**Figure 4.**
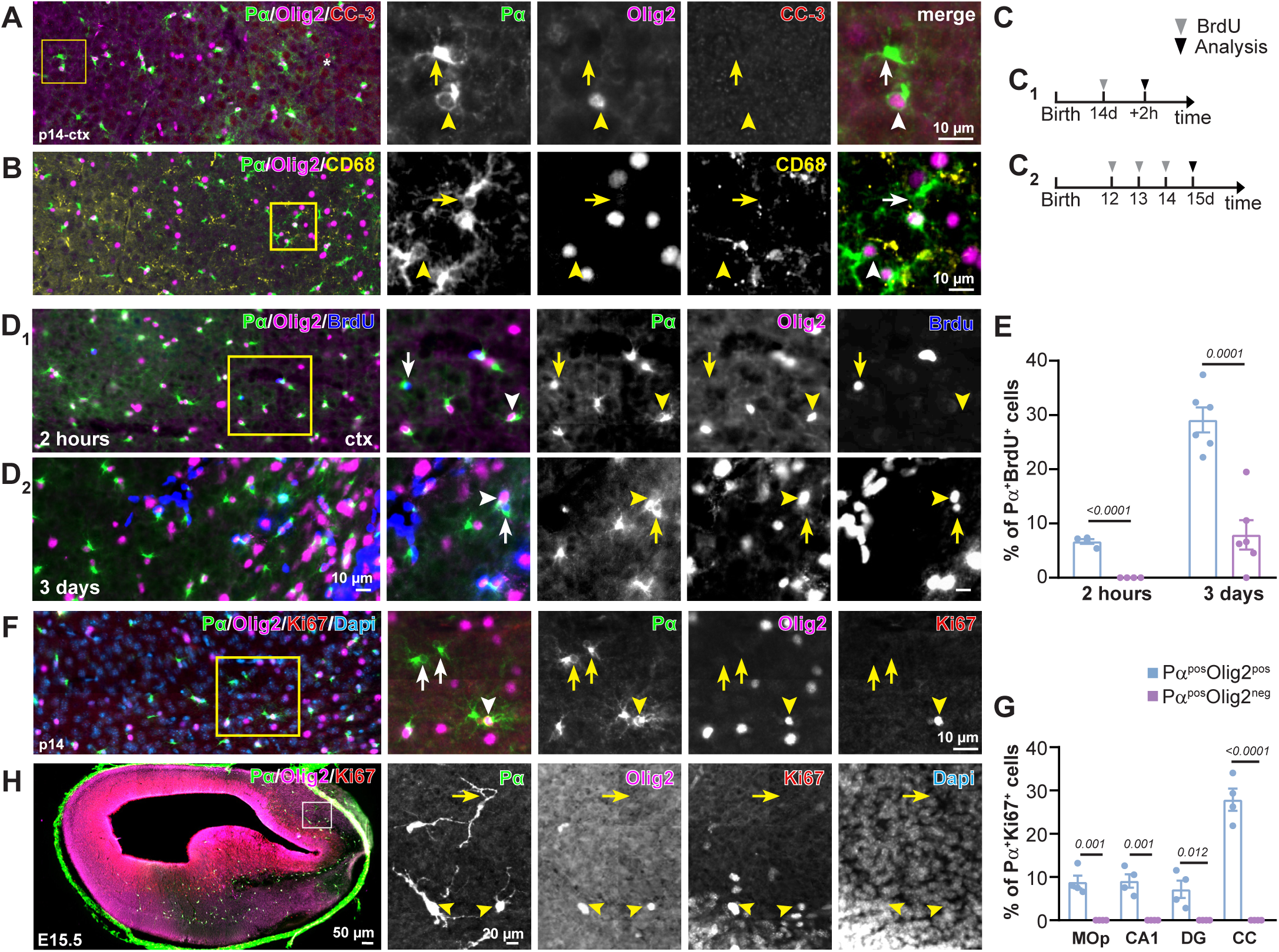
Olig2^neg^ cells proliferate less than Olig2^pos^ OPCs. **A**, Pα^pos^Olig2^neg^ cells lacked immunolabeling by apoptotic markers. **B**, Pα^pos^Olig2^neg^ cells are not phagocytosed by microglia. Arrowheads: Olig2^pos^ OPCs, arrows: Olig2^neg^ OPCs. **C**, Experimental schedule for BrdU assay (**C_1_** and **C_2_**) in **D_1_** and **D_2_**, respectively. **D,** Immunostaining of Pα and Olig2 in the cortex of p14 mice after 2 hours (**C_1_, D_1_)** or 3 days (**C_2_, D_2_**) of BrdU administration. **E**, Quantification of proliferating Olig2^pos^ and Olig2^neg^ OPCs by BrdU labeling. **F, G,** Unlike Olig2^pos^ OPCs, Olig2^neg^ OPCs did not express the proliferation marker Ki67 (primary motor cortex, MOp; CA1; dentate gyrus, DG; corpus callosum, CC). **H,** Pα^pos^Olig2^neg^ cells did not exhibit Ki67 immunoreactivity in the forebrain at embryonic day 14.5. Arrowheads: Olig2^pos^ OPCs, arrows: Olig2^neg^ OPCs.

Then, we assessed the proliferation by BrdU incorporation. To distinguish the fast and slowly diving cells, BrdU was administrated at 2 hours prior to the analysis at p14 or for consecutive 3 days from p12 till p15 (Fig. 4C). With the short pulse of BrdU administration, none of the Olig2^neg^ OPCs showed BrdU immunoreactivity in contrast to about 7 % of Olig2^pos^ OPCs (Fig. 4D, E). Nevertheless, a long-term administration of BrdU labeled about 10 % of Olig2^neg^ OPCs with BrdU, which was still less than Olig2^pos^ OPCs (29 %) (Fig. 4D, E), suggesting that Olig2^neg^ OPCs divide very slowly if at all, and are likely derived from Olig2^pos^ OPCs.

To further clarify a potential proliferation capacity of Olig2^neg^ cells, we performed an immunostaining against the mitotic marker Ki67 on these cells at p14 brain (Fig. 4F). About 7-9 % of Olig2^pos^ OPCs in the grey matter (MOp, CA1 and DG) and about 28 % in the white matter (corpus callosum) (Fig. 4G) (arrowheads in Fig. 4F) were positive for Ki67, whereas none of the Pα^pos^Olig2^neg^ cells showed immunoreactivity to Ki67 in any of the brain regions (Fig. 4G, Supplementary Fig. 4A). To substantiate that Olig2^neg^ cells are not proliferative, we applied the same immunohistochemical protocol in the forebrain of embryos at embryonic day (E) 15.5, when the cells are more actively dividing. At E15.5, the first and second wave of OPCs are already generated (Kessaris et al., 2006). We still could detect about 7 % of OPCs being Olig2^neg^ (Fig. 4H), suggesting Olig2^neg^ is not limited to the third wave of OPCs generated at perinatal days. While the majority of Olig2^pos^ OPCs expressed Ki67, none of the Olig2^neg^ cells expressed Ki67, thereby strongly suggesting a non-proliferative property of Olig2^neg^ OPCs.

Taken together, our data suggest that Olig2^neg^ cells persist in the adult brain as a distinct subpopulation of oligodendrocyte lineage cells derived from Olig2^pos^ OPCs.

### Olig2^neg^ cells exhibit a simplified morphology

Since Olig2 is critical for OPC differentiation and subsequent myelination, we asked whether these Olig2^neg^ cells still could generate oligodendrocytes. OPCs increase branching of processes as they differentiate into oligodendrocytes (Pfeiffer, Warrington, & Bansal, 1993). Therefore, we compared the morphology of these cells from the p14 cortex using a Sholl analysis based on Pα immunostaining (Fig. 5A). We found that Olig2^neg^ OPCs displayed less process branches and shorter total filament length compared to the Olig2^pos^ cells (Fig. 5B-D). These data indicate that Olig2^neg^ cells might remain in a precursor stage rather than differentiating into oligodendrocytes.

**Figure 5.**
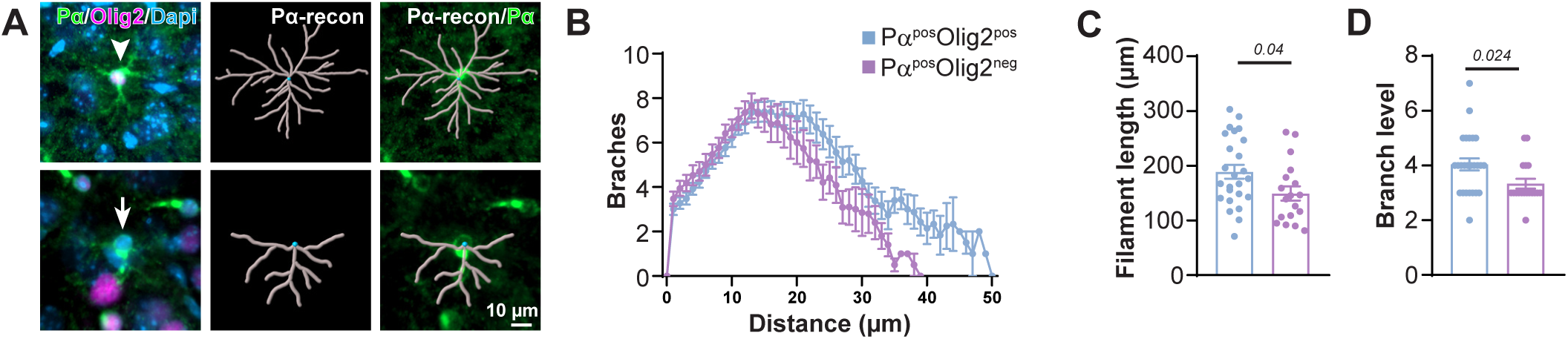
Pα^pos^Olig2^neg^ cells are morphologically less complex than Olig2^pos^ OPCs. **A**, Morphological analysis of Olig2^pos^ and Olig2^neg^ OPCs based on Pα immunostaining using Imaris software. **B-D**, Sholl analysis showed shorter filament length (**B, C**) and less branches (**B, D**) of Olig2^neg^ OPCs.

Studies have suggested that perivascular OPCs exhibit simpler morphologies in comparison to parenchymal ones (Kishida et al., 2019). To understand whether these Olig2^neg^ cells are perivascular OPCs, we labelled endothelial cells of the brain vasculature by PECAM-1/CD31 immunostaining in the p14 brain (Fig. 6A, Supplementary Fig. 4B). About 40 % of Olig2^neg^ OPCs in CC and 30 % in CA1 were located at blood vessels (Fig. 6B). However, we also observed a rather low percentage of total Olig2^pos^ OPCs situated perivascular (18.5 ± 8.2 % in CC; 16.7 ± 2.7 % in CA1). In addition, in cortex and DG, the percentages of these two subtypes at blood vessels were similar (27.7 ± 8.4 % (Olig2^neg^) vs 25.9 ± 7.0 % (Olig2^pos^) in MOp; 21.3 ± 2.1 % (Olig2^neg^) vs 15.9 ± 2.8 % (Olig2^pos^) in DG) (Fig. 6B). Hence, our results indicate that Olig2^neg^ cells are not necessarily perivascular OPCs, and vice versa. Interestingly, at E15.5 forebrain (about 7 % of OPCs were Olig2^neg^), almost 80 % of Olig2^neg^ cells were situated perivascular (Fig. 6C-E). This number was 2-3 folds higher than those observed in the adult brain (Fig. 6B).

**Figure 6.**
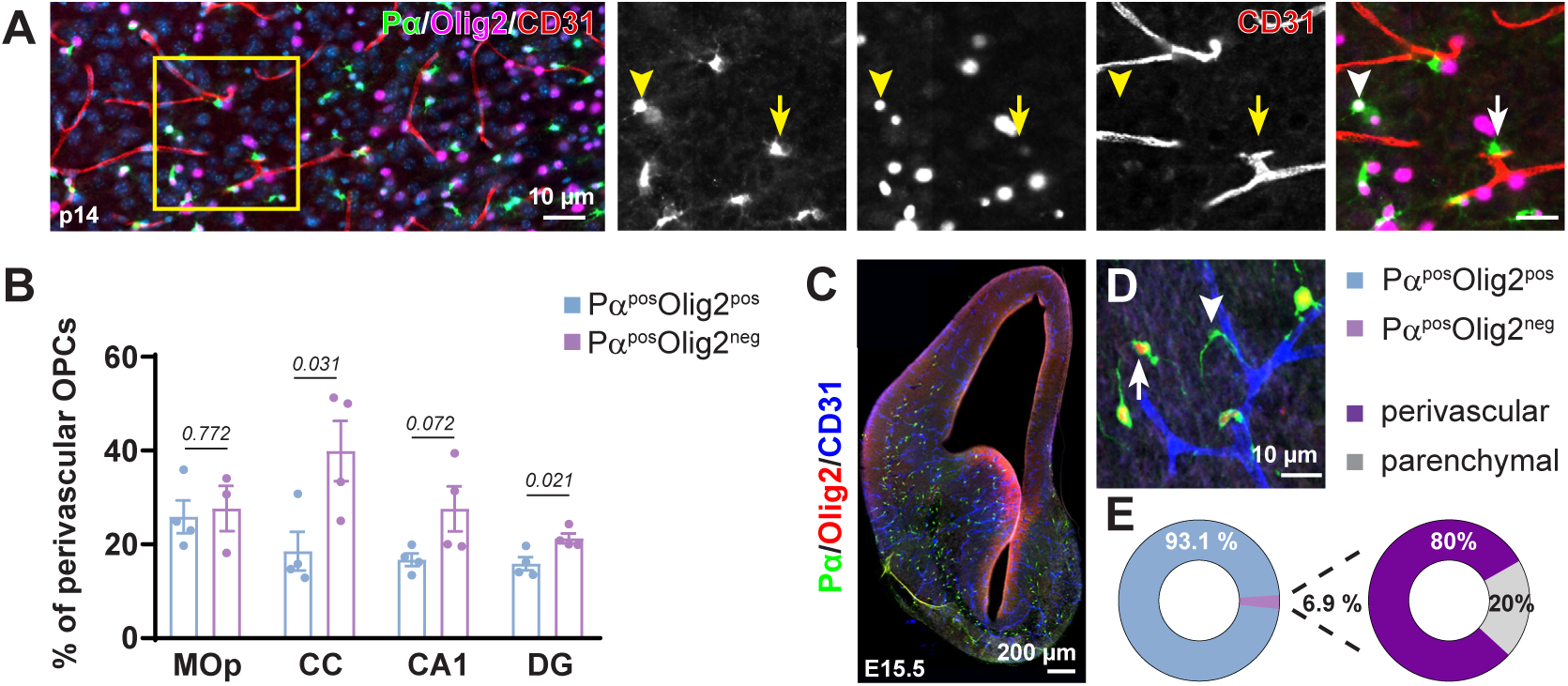
In dentate gyrus and corpus callosum Pα^pos^Olig2^neg^ cells are preferentially located along the blood vessels in comparison to Olig2^pos^ OPCs. **A**, Immunostaining of Pα, Olig2 and CD31 at the p14 cortex showed OPCs with or without Olig2 expression adjacent to the blood vessels. **B**, Percentage of perivascular OPCs with or without Olig2 from total Pα^pos^Olig2^pos^ or Pα^pos^Olig2^neg^ cells in p14 brain, respectively. **C, D**, Immunostaining of Pα, Olig2 and CD31 in the E15.5 forebrain. **E**, Population of Olig2^neg^ OPCs among all OPCs and the percentage of perivascular Olig2^neg^ OPCs from all Olig2^neg^ OPCs at E15.5. Arrowheads: Olig2^pos^ OPCs, arrows: Olig2^neg^ OPCs.

These results indicate that OPCs can down regulate Olig2 and stay in their precursor phase during brain development.

### Increase of Olig2^neg^ OPCs in the adult brain after acute brain injuries

The main establishment of neural networks occurs between birth and p30, which parallels with the emergence of Olig2^neg^ OPCs. OPCs shape neural circuits by participating in synaptic transmission already from p5 and forming connections after differentiation (Bergles, Roberts, Somogyi, & Jahr, 2000; Lin & Bergles, 2004; Orduz et al., 2015). Therefore, we asked whether there could be a causal link between the brain activity status and the formation of Olig2^neg^ OPCs. Therefore, we performed three different types of acute brain injuries in adult mice, when the Olig2^neg^ cell population is rather low. We used stab wound injuries (SWI), kainic acid (KA)-evoked seizures or middle cerebral artery occlusion (MCAO). SWI was induced to 9-week-old mice and analyzed at 3, 7 or 10 days post injury (Fig. 7A, B). Already at 3 days post injury (dpi), the density of Pα^pos^Olig2^neg^ cells in the ipsilateral side, especially at the lesion site (50 µm aside from the lesion), increased 10 fold compared to the contralateral side (34.1 ± 6 vs 3.5 ± 1 cells/1x10^-2^ mm^3^, Fig. 7C). This number increased further at 7 dpi (75.8 ± 17.5 cells/1x10^-2^ mm^3^), while returned to the level as 3 dpi at 10 dpi (28.6 ± 1.8 cells/1x10^-2^ mm^3^) (Fig. 7C). KA injection and MCAO also triggered the increase of Pα^pos^Olig2^neg^ OPC population at the ipsilateral cortex 2 wp KA injection (Fig. 7D-F) or in the most affected region (according to the GFAP and Iba1 expression) of the MCAO stroke model (Supplementary Fig. 5). Further immunostaining showed that these Olig2^neg^ OPCs did not express Ki67 (Fig. 8A), as observed under physiological conditions. All these data indicate that the formation of Olig2^neg^ cells can be triggered in the adult brain by modifying the brain activity.

**Figure 7.**
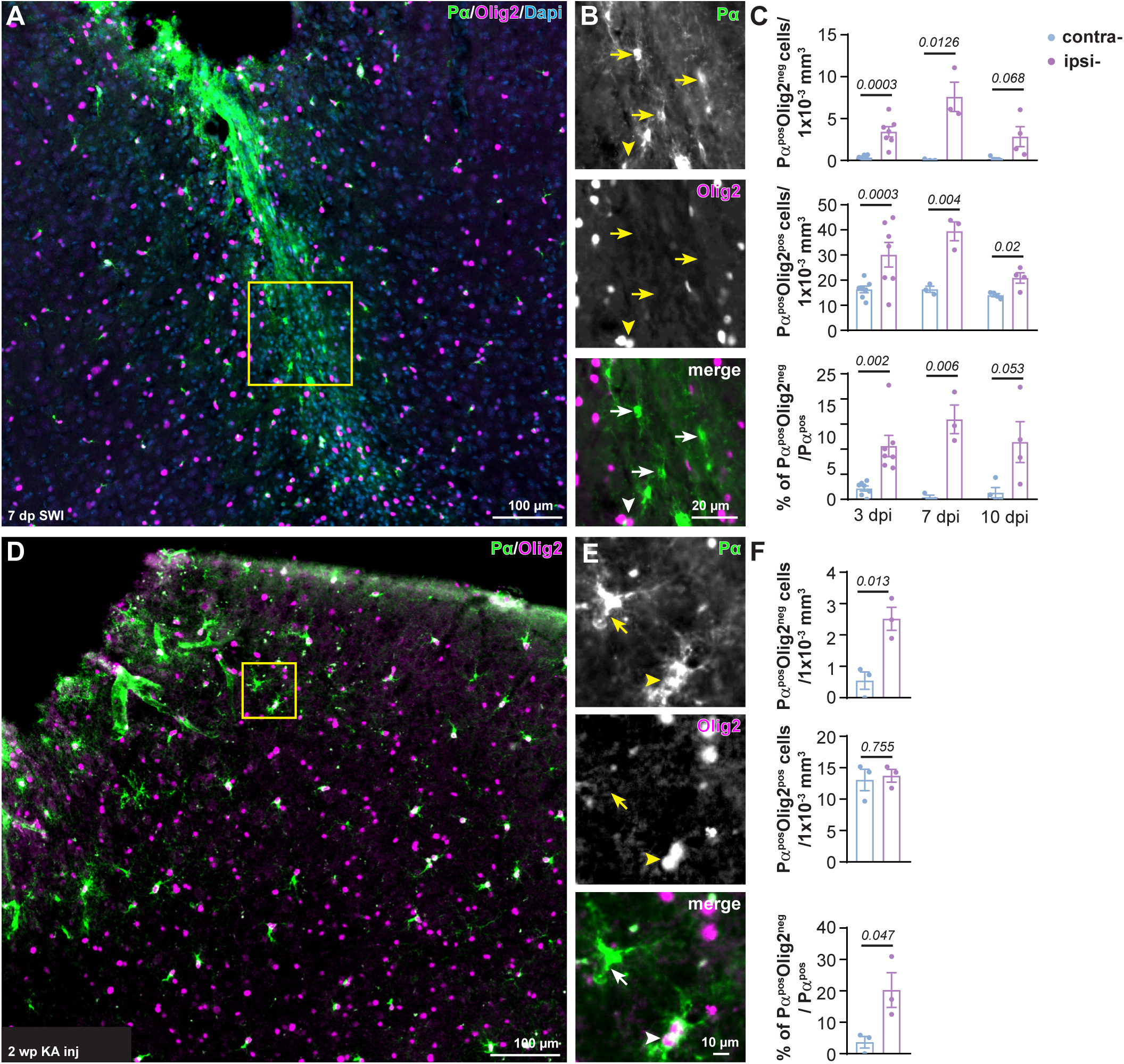
Cortical brain injuries trigger the new formation of Pα^pos^Olig2^neg^ OPCs at the lesion site. **A**, **B**, Immunostaining of Pα and Olig2 at the ipsilateral side after 7 days of stab wound injury (SWI) shows numerous Pα^pos^Olig2^neg^ OPCs at the lesion site. **C**, Quantification of Olig2^neg^ and Olig2^pos^ OPCS cell density, as well as the proportion of Pα^pos^Olig2^neg^ cells among all OPCs in the contralateral and ipsilateral side at 3, 7, and 10 days post SWI. **D, E**, Immunostaining of Pα and Olig2 at the ipsilateral side of kainate injected cortex 2wpi. **F**, Cell densities of Olig2^neg^ and Olig2^pos^ OPCs, and the percentage of Olig2^neg^ cells in the contralateral and ipsilateral side at 2 wpi of kainate. Arrowheads: Olig2^pos^ OPCs, arrows: Olig2^neg^ OPCs.

**Figure 8.**
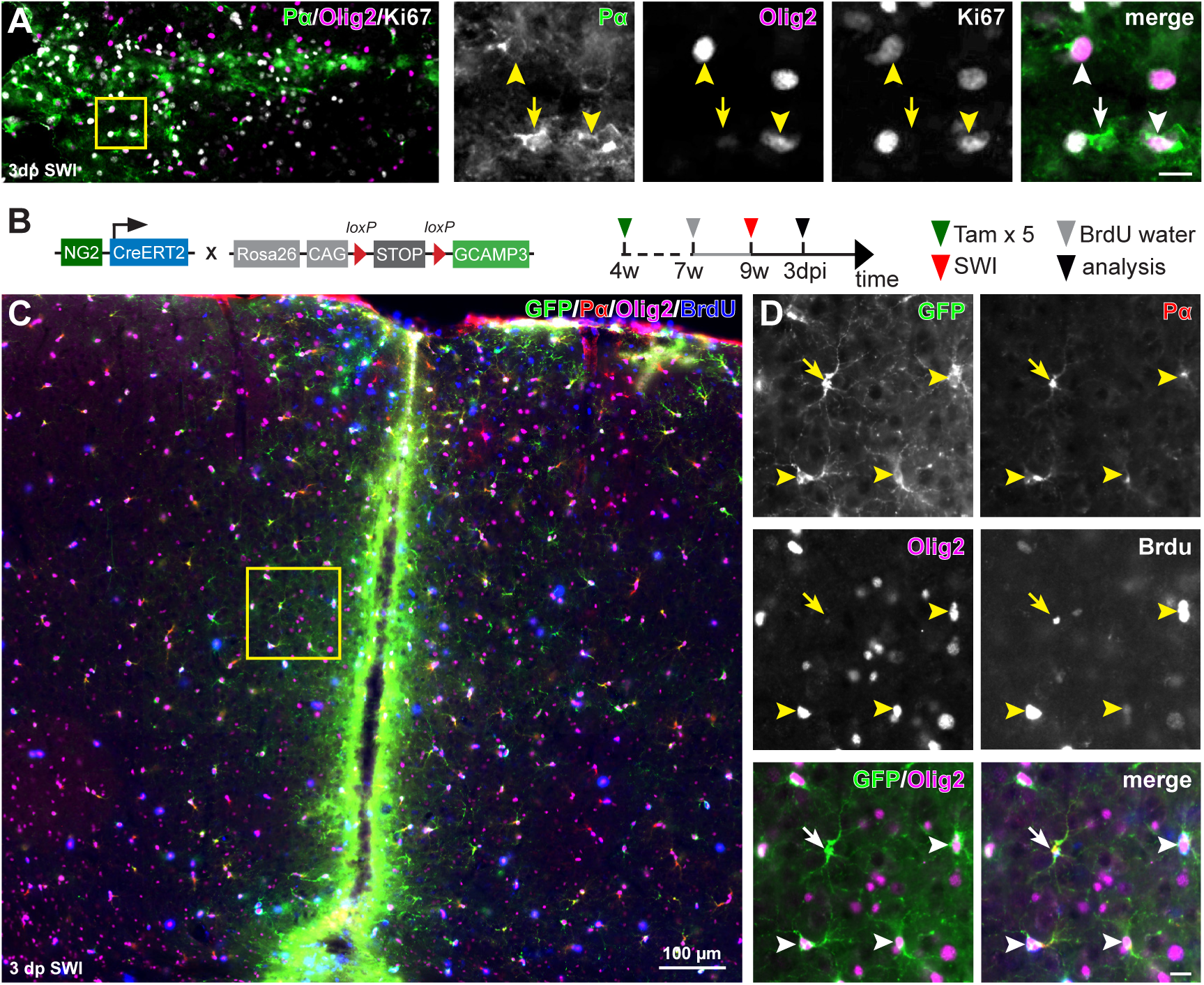
OPCs transiently suppress Olig2 after acute brain injury. **A,** Immunostaining of Pα^pos^Olig2^neg^ OPCs by the mitotic marker Ki67 in the lesion site at 3 days post stab wound injury (dpi SWI). **B**, Scheme of transgene structure and experimental design. **C**, Immunostaining of GFP, Pα, Olig2 and BrdU at the lesion site 3dpi. **D**, Representative images showing that GFP^pos^Pα^pos^ OPCs with (arrowhead) or without Olig2 (arrow) expression incorporated with BrdU. Arrowheads: Olig2^pos^ OPCs, arrows: Olig2^neg^ OPCs.

To investigate whether these Pα^pos^Olig2^neg^ cells were derived from pre-existing OPCs or from other precursors, we took advantage of NG2-CreER^T2^ x R26-lsl-GCaMP3 mice where the OPCs can be assessed by evaluating the reporter gene GCaMP3 using GFP antibodies. Recobination was induced by injection of tamoxifen at the age of four weeks. An SWI was performed at the age of nine weeks (Fig. 8B). About 90 % OPCs were recombined at the age of 9 weeks (Fang et al., 2022). Analyzed at 3 dpi, all the Pα^pos^Olig2^neg^ cells in the ipsilateral side were GFP^pos^ (Fig. 8C, D), indicating that Pα^pos^Olig2^neg^ cells were originating from pre-existing GFP^pos^ OPCs, not from other precursor cells. To confirm these hypothesis, we administered BrdU in the drinking water of NG2- CreER^T2^ x R26-lsl-GCaMP3 mice for two consecutive weeks prior to the SWI to label dividing OPCs (Fig. 8B). In the contralateral side of 3 dpi cortex, about 47.5 % of Olig2^pos^ OPCs were BrdU^+^, but none of the Olig2^neg^ cells, again excluding a proliferative capacity of Olig2^neg^ cells. However, at the lesion site, 7 % of Pα^pos^GFP^pos^Olig2^neg^ OPCs had incorporated BrdU (Fig. 8C, D), suggesting their origin from pre-existing OPCs after acute brain injuries.

Taken together, our data demonstrate that acute brain injuries can induce OPCs to stop Olig2 expression.

### The formation of Olig2^neg^ OPCs is related to the establishment of neural networks

To substantiate that Olig2^neg^ OPCs are required for the plastic formation of functional neural networks, we challenged mice with a three-week complex running program on the Erasmus Ladder (Van Der Giessen et al., 2008). After five days of habituation, mice at 8 or 19 weeks of age had to run the ladder with random cues and obstacles for consecutive 16 days (Fig. 9A). To correlate the learning and the appearance of Olig2^neg^ OPCs, immunostaining was performed at three different time points: before learning (right after habituation, day 5), middle of learning (day 13) and end of learning (day 21) (Fig. 9A, Supplementary Fig. 6A). Quantification of missteps showed that the mice could run properly only at the last few days of the learning (Supplementary Fig. 6B). Learning a complex running program combines basic motor activity and learning. Therefore, motor cortex and hippocampus were primarily analyzed for the Pa^pos^Olig2^neg^ cell population, both brain regions closely related to motor activity and learning in the forebrain (Burman, 2019; Jacobacci et al., 2020). In the hippocampus, while the density of Pa^pos^Olig2^neg^ cells increased 2-4 times in both DG and CA1 region compared to control animal after complete learning session (Fig. 9A-D), no change was observed before or during the early learning period (Supplementary Fig. 6C-E). In addition, the population of Olig2^neg^ OPCs remained stable in the motor cortex, somatosensory cortex or CC (Supplementary Fig. 6F-J). These results suggest that the formation of Olig2^neg^ OPCs in the hippocampus is likely related to learning induced novel neural networks.

**Figure 9.**
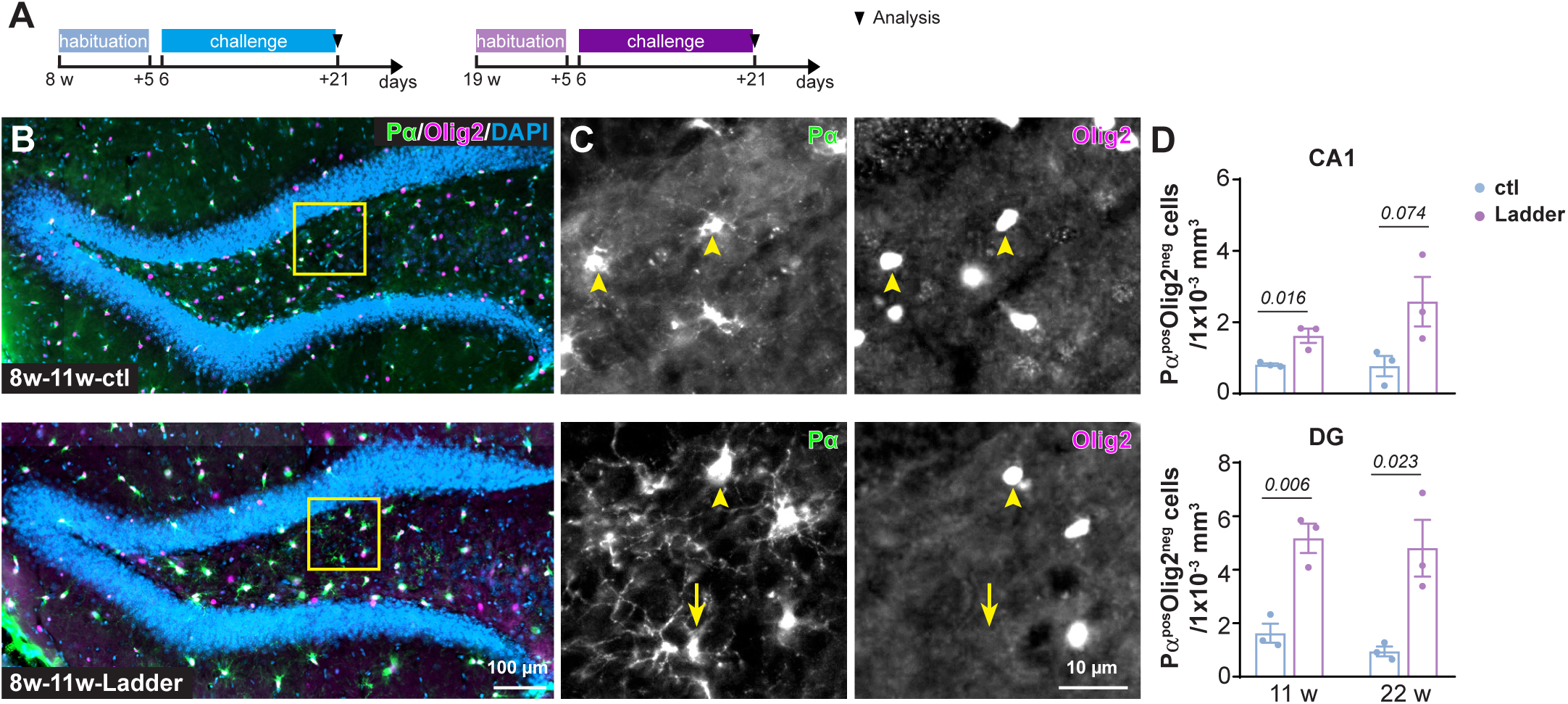
Complex motor learning increases the formation of Pα^pos^Olig2^neg^ OPCs in the adult hippocampus. **A**, Experimental schedule. **B,** Overview of hippocampus immunostained with Pα and Olig2 in the control and trained mice. **C**, Magnified views of the area indicated in **B** show OPCs (Pα^pos^) with (arrowhead) or without (arrow) Olig2 expression. **D**, Quantification of Pα^pos^Olig2^neg^ cell density in the CA1 and DG region in the control and Erasmus Ladder trained mice at 11 week or 22 week. Arrowheads: Olig2^pos^ OPCs, arrows: Olig2^neg^ OPCs.

To further understand the biological meaning of Olig2 silencing in OPCs, we compared the genes differentially expressed by Olig2^neg^ and Olig2^pos^ OPCs using the published single cell transcriptomic data (Marques et al., 2016) (https://mouse-oligo-het.cells.ucsc.edu). Firstly, we assessed whether Olig2^neg^ OPCs form any clusters in UMAP dimension analysis. Defined by the expression of *Pdgfra* and non-expression of *Plp1* from different brain regions of PDGFRα-Cre x R26-CAG-EGFP^fl/fl^ mice at different ages, Olig2^neg^ OPCs did not exhibit cluster in the whole OPC population (Fig. 10A). These data suggest that Olig2^neg^ OPCs are rather generally present in all ages and brain regions. Then we further compared the genes differentially expressed in these two populations and observed 59 genes enriched in Olig2^neg^ OPCs and 400 for Olig2^pos^ cells (Supplementary File 1), and top 20 genes for each population are listed in Fig. 10B. In Olig2^neg^ cells, for example, adenosine A1 receptor (A1AR, *Adora1*) was enriched compared to Olig2^pos^ cells, suggesting a potential purinergic signaling involved in suppression of Olig2 in OPCs. Shown by gene ontology (GO) analysis, genes positively regulating cell differentiation and myelination, eg. *Olig2, Mbp, Egr2* were highly enriched in the Olig2^pos^ cells (Fig. 10C).

**Figure 10.**
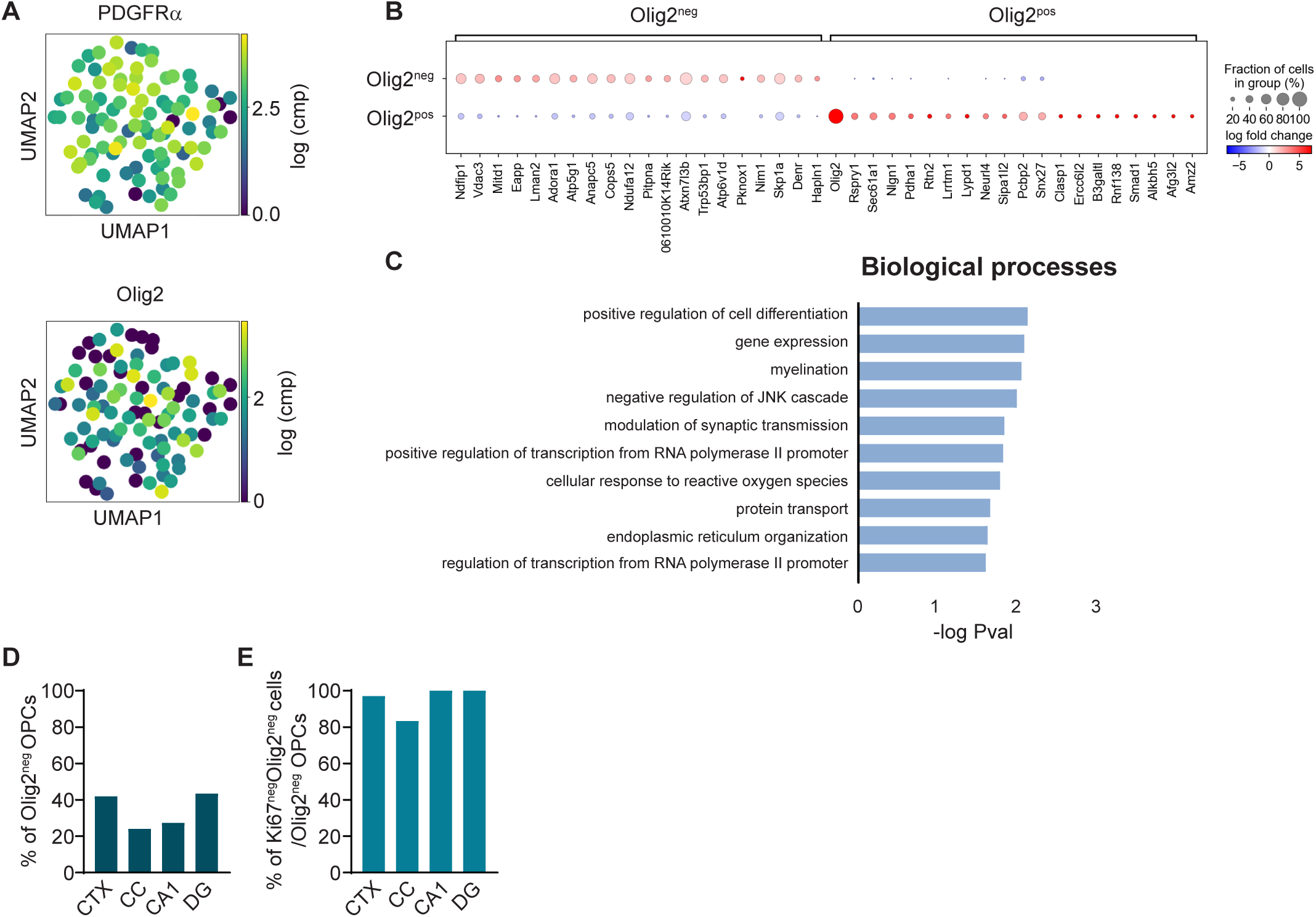
Cell differentiation and myelination is suppressed in Olig2^neg^ OPCs. **A**, UMAP dimension reduction representation of single OPCs for PDGFRα and Olig2 expression from published database (Marques et al., 2016). **B,** Dot plot visualizing differential gene expression between Olig2^neg^ and Olig2^pos^ OPCs. **C,** Biological processes enriched for genes that are higher expressed in Olig2^pos^ OPCs. **D, E**, Quantification of OPCs expressing no Olig2 (expression level=0) (**D**) and Olig2^neg^ OPCs expressing no Ki67 (**E**) in cortex (CTX), corpus callosum (CC), CA1 and dentate gyrus (DG) from the data source of (Marques et al., 2016).

In summary, our results indicate that a subset of OPCs transiently downregulate Olig2 upon micro-environmental changes, contributing further to OPCs heterogeneity.

## Discussion

Olig2, a basic helix loop helix transcription factor, is a critical determinant for oligodendrocyte lineage function, specification and differentiation. However, here, we observed a subset of OPCs that lack Olig2 throughout life in the brain, in grey as well as in white matter regions such as cortex, hippocampus or corpus callosum. Emergence of this population coincided with brain activity changes. Unlike Olig2^pos^ OPCs, Olig2^neg^ cells exhibited very low (if any) self-renewing activity and a simplified morphology in the developing brain, suggesting a putative functional difference between these two subpopulations. Overall, our study demonstrated that upon the change of brain activity, a subset of OPCs transiently shutdown Olig2 expression.

OPCs receive glutamatergic and GABAergic input from neurons and differentiate into oligodendrocytes (Bergles et al., 2000; Gautier et al., 2015; Orduz et al., 2015; Serrano-Regal et al., 2020). Not only, OPCs also shape neural circuits (Fang et al., 2022; Xiao, Petrucco, Hoodless, Portugues, & Czopka, 2022; X. Zhang et al., 2021). For instance, prior to myelin onset, the large population of OPCs in mouse medial prefrontal cortex release TNF like weak inducer of apoptosis (TWEAK) to optimize interneuron population which is pivotal for proper neural network activity (Fang et al., 2022). In addition, as seen in zebrafish optic tectum, a subtype of OPCs remain precursors throughout life and control axonal remodeling, never differentiating into myelinating oligodendrocytes (Xiao et al., 2022). A recent study has demonstrated that a set of genes, involved in pathways including neuronal differentiation and brain development, are repressed by Olig2 to ensure OPC differentiation into oligodendrocytes (K. Zhang et al., 2022). Therefore, it is tempting to speculate that OPCs switch off Olig2 expression and remain as precursors or NG2 glia upon the change of brain activity. Indeed, we observed a less complex morphology of Olig2^neg^ cells in comparison to Olig2^pos^ cells, suggesting Olig2^neg^ OPCs yet exhibit less potential to commit the differentiation.

In general, neural plasticity is greater in the developing brain than in the adult, and learning a new skill or brain injuries could boost neural circuit reorganization (Galván, 2010; Su, Veeravagu, & Gerald, 2016). The appearance of Olig2^neg^ OPCs in the postnatal brain coincided with the development of neuron-OPCs connectivity starting from postnatal day 4-5 and reaching a peak at p10 for cortical interneurons (Orduz et al., 2015). Complex motor learning processes as well as acute brain injuries evoked novel formation of Olig2^neg^ OPCs. Interestingly, BMP4, known to repress Olig2 expression, is essential for synapse plasticity and is upregulated during brain development and after injuries (Bai et al., 2021; Higashi, Tanaka, Iida, & Okabe, 2018). Therefore, BMP4 might be involved in the downregulation of Olig2 in OPCs during neural circuit establishment and to keep OPCs in precursor stage and innervate into neural circuit. Nevertheless, Olig2^neg^ cells were more frequently observed in white matter areas such as corpus callosum than in the grey matter of cortex or hippocampus. This could be attributable to inherited intrinsic programs (Viganò & Dimou, 2015; Viganò et al., 2013) and/or microenvironmental variations. E.g., the corpus callosum exhibits higher stiffness than grey matter, and such high stiffness hinders OPCs proliferation and differentiation (Segel et al., 2019). However, the Olig2^neg^ OPC population wanes with age, albeit the brain stiffness increases with age (Segel et al., 2019). Hence, molecular mechanisms involved in the generation of Olig2^neg^ OPCs needs to be further studied.

Taking advantage of NG2-CreER^T2^ mice, we found that Olig2^neg^ OPCs were derived from pre-existing OPCs downregulating Olig2 upon acute brain injuries. Interestingly, different to the Olig2^pos^ cells, Olig2^neg^ OPCs were rarely recognized by Ki67 antibody at any brain region. Indeed, Olig2 directly activates cell proliferating pathways to facilitate tumor growth in proneural glioma (F. Lu et al., 2016). Single cell RNA sequencing database of zebrafish and mouse brain have also suggested a subgroup of PDGFRα^pos^ OPCs that apparently lack expression of the mitotic marker Ki67 (Marisca et al., 2020; Marques et al., 2016). By analyzing the published database (https://mouse-oligo-het.cells.ucsc.edu) (Marques et al., 2016), we identified slightly higher numbers of OPCs without Olig2 mRNA (31.6 % (database) vs 10-25% (own data), Fig. 10D) and also similarly large proportion of these cells not expressing Ki67 (91 % (database) vs 100% (own data), Fig. 10E). In addition, in the zebra fish spinal cord, the ‘cluster #1’ OPCs were also found to be negative for Ki67, considered as ‘quiescent’ OPCs since they lacked proliferation and differentiation related markers (Marisca et al., 2020). Instead, these cells were enriched with mRNAs involved in axon guidance and synaptic communication (Marisca et al., 2020). Hence, if Olig2^neg^ OPCs represent similar ‘quiescent’ OPC population in the mouse brain, these cells might be also well integrated into neural circuit. Nevertheless, whether Olig2^neg^ OPCs are a functionally and physiologically different group of OPCs needs additional further characterization, e.g. by electrophysiology or patch-sequencing.

Developmentally, OPCs are generated in three waves: at E12.5 from Nkx2.1^pos^ precursors, at E14.5 from Gsx2^pos^ precursors and from Emx1^pos^ precursors at perinatal days (Kessaris et al., 2006). After birth, the first wave of OPCs, especially in the dorsal cortex, disappear within the first two postnatal weeks, matching to the time points when Olig2^neg^ OPCs start to appear. However, we did not observe Olig2^neg^ OPCs expressing cleaved caspase-3 or being phagocytosed by microglia. In addition, Olig2^neg^ OPCs can also exist in the embryonic brain (detected at E14.5), when yet no OPC death was reported. Nonetheless, we do not exclude other mechanisms eliminating OPCs thereby transiently becoming PDGFRα^pos^Olig2^neg^ cells, when PDGFRα is yet not fully degraded whereas Olig2 already is. However, the half-life of PDGFRα is 3 hours in the absence of ligands and that of Olig2 is in the range of 4-8 hours, at least *in vitro* (Coats, Olashaw, & Pledger, 1994; Kupp et al., 2016). Unfortunately, these Olig2^neg^ OPCs are not able to be fate-tracked by live imaging techniques, since Olig2^neg^ OPCs can, so far, only be recognized by the immunostaining.

At the early postnatal brain Olig2 is mainly expressed by oligodendrocyte lineage cells and a subtype of astrocytes (Cai et al., 2007; Wang et al., 2021). Subsequently, astrocytes progressively downregulate Olig2 during their postnatal development, thereby Olig2 becomes restricted to the oligodendrocyte lineage (Q. R. Lu et al., 2000; Takebayashi, Nabeshima, Yoshida, Chisaka, & Ikenaka, 2002; Zhou et al., 2000). However, under pathological conditions, Olig2 expression alters and the expressing population even extends to other cell types, e.g. astrocytes (Chen et al., 2008). Olig2 is essential for the astrogliosis after cortical injuries (Chen et al., 2008). Deletion of Olig2 specifically in OPCs induced the differentiation into astrocytes rather than into oligodendrocytes (Zhu et al., 2012). Several studies have indicated that OPCs differentiate into astrocytes upon acute brain injuries (Bai et al., 2021; Dimou, Simon, Kirchhoff, Takebayashi, & Goetz, 2008; Komitova, Serwanski, Lu, & Nishiyama, 2011; Scheller, Bai, & Kirchhoff, 2017; Tatsumi et al., 2008). Therefore, it is possible that these Olig2^neg^ OPCs switch their fate and give rise to astrocytes. However, fate mapping with NG2-CreER^T2^ mice did not show any reporter positive astrocytes in the physiological brain, also not after the peak of Olig2^neg^ OPCs appearance. Therefore, additional studies are necessary to investigate the function of Olig2^neg^ OPCs. This might provide a novel insight for the understanding of neuron-OPCs communication under physiological and pathological conditions.

## Materials and Methods

### Ethics statement

All animal experiments were carried out at the University of Saarland in strict accordance with recommendations of European and German guidelines for the welfare of experimental animals. Animal experiments were approved by Saarland state’s “Landesamt für Gesundheit und Verbraucherschutz” in Saarbrücken/Germany (animal license numbers: 36/2016, 03/2021, 08/2021).

### Animals

All mouse lines were maintained in C57BL/6N background and housed with a 12 hour (h) light/dark cycle at 20°C in the animal facility of the CIPMM. Mice were fed a breeding diet (V1125, Sniff) *ad libitum*. For the developmental study, C57BL/6N animals were analyzed at the age of postnatal day 5 (P5), P9, two weeks (2w), 4w, 9w, 11w, 22w, 44w, while embryos were analyzed at E15.5. To follow the fate of OPCs, we took advantage of NG2-CreERT2 knock-in mice carrying CAG-^fl^STOP^fl^-tdTomato (TgH(ROSA26-CAG-fl-stop-fl-tdTomato)) or CAG-^fl^STOP^fl^-GCaMP3 (TgH(ROSA26-CAG-fl-stop-fl-GCaMP3)) reporter (W. Huang et al., 2014; Madisen et al., 2010; Paukert et al., 2014). Mice were always heterozygous for NG2-CreER^T2^ and homozygous for the floxed reporter loci.

### Tamoxifen induced recombination

Tamoxifen (Carbolution, Neunkirchen, Germany) was dissolved in Miglyol®812 (Caesar & Lorentz GmbH, Hilden, Germany) to a final concentration of 10 mg/ml and administrated intraperitoneally for two consecutive days at postnatal day 7 and 8 or for five consecutive days at the age of 4 weeks (Fang et al., 2022; Jahn et al., 2018).

### Erasmus Ladder

Complex motor learning was performed using the Erasmus Ladder (Noldus Technology) as previously described with modifications (Saab et al., 2012; Van Der Giessen et al., 2008). Individual test mice were placed in a dark shelter. With a light cue and 3 second delayed air cue, the animal was encouraged to run over the ladder and reach the shelter at the other side. One trial was defined as a single ladder crossing from one shelter to another, and one session was composed of 42 trails. Each mouse performed one session per day. The first five days were regarded as training sessions followed by sixteen days of challenging sessions. At training sessions, mice were habituated with the ladder and the cues. From the sixth day, during ladder running mice were randomly perturbed by single sounds or obstacles, or combinations of sound and obstacles. Only one or no perturbation was added per trial. The duration time of steps per trial was automatically recorded.

### Stab wound injury

Stab wound injuries (SWI) were performed on 9-12 weeks old mice as described previously (Bai et al., 2021). Briefly, mice were anesthetized with a mixture of 2 % Isoflurane and 49 % O_2_ and 49 % N_2_O via inhalation and the skull was thinned with a dental drill laterally 1.5 mm and longitudinally 2 mm from Bregma. A 1 mm deep stab wound was made by the insertion of surgical scalpel into the somatosensory cortex. The skin was sutured and buprenorphine was intraperitoneally given daily for three consecutive days. Tramadol (0.4mg/ml) was administrated in drinking water for 7 consecutive days. Mice were analyzed at 3, 7 and 10 days post injury (dpi).

### Middle cerebral artery occlusion

Middle cerebral artery occlusion (MCAO) was performed as previously described (W. Huang et al., 2020). Mice were anesthetized with inhalation anesthetics as described above. Mice were anesthetized with inhalation anesthetics as described above. First, the left common carotid artery (CCA) and the external carotid artery were permanently ligated with silk sutures. After performing an arteriotomy on the CCA, a silicon-coated filament (Doccol Corp, CA) was inserted into the CCA and advanced through the internal carotid artery until it reached the origin of the middle cerebral artery. After 15 minutes of occlusion, reperfusion was obtained by withdrawal of the filament. Lastly, another suture was made around the CCA, to prevent back flow through the arteriotomy.

Mouse body temperature was continuously monitored using a rectal thermometer and an adjustable heat plate. Mice received intraperitoneal injection of buprenorphine for pain relief and subcutaneous injection of 0.5 ml saline as fluid replacement for three consecutive days. Tramadol (0.4mg/ml) was administrated in drinking water for 7 consecutive days.Mice were analyzed at 3 dpi.

### Kainate injection

Cortical kainate injections were performed unilaterally as described previously (Bedner et al., 2015). Briefly, 70 nl of 20 mM kainic acid in saline was injected to the right hemisphere of anesthetized mice, at the position of AP:-1.92 mm, ML: 1.5 mm, DV: 1 mm referred to the bregma. The skin was sutured and buprenorphine was intraperitoneally given daily for three consecutive days. Tramadol (0.4mg/ml) was administrated in drinking water for 7 consecutive days. Mice were analyzed at 1 and 14 dpi.

### Immunohistochemistry

Mice were perfused with PBS followed by 4 % PFA. After post fixation, coronal vibratome slices were collected. After 1 h incubation with blocking buffer (5 % horse serum with 0.5 % Triton in PBS), free floating slices were incubated with primary antibodies at 4 °C overnight, followed by secondary antibody incubation. DAPI (25 ng/ml) was used to stain nuclei (A10010010, Biochimica). Primary and secondary antibodies are listed in the **Supplementary Table 1** and **2**, respectively.

### Bromodeoxyuridine assay

For adult animals, 1 mg/ml BrdU was administered in the drinking water for two weeks before the surgery. For short pulse labeling, 10 mg/ml BrdU saline solution was intraperitoneally injected to the mice 2 h prior to the analysis. For BrdU immunostaining, after washing off the secondary antibody, slices were fixed with 2 % PFA at room temperature (RT) for 15 min, followed by washing with 1x PBS twice and once by distilled water. Slices were then incubated with 2 M HCl at 37°C for 45 min. After three times washing, slices were incubated with primary BrdU antibody and secondary antibodies as described above.

### Microscopic analysis and quantification

Two brain slices per mouse and at least three animals per group were analyzed. Images were obtained with an automated slide-scanning epifluorescence microscopy system (Zeiss AxioScan Z1, for overviews) or a confocal laser-scanning microscope (Zeiss LSM-710, for high-resolution and image stacks).

For quantification of fluorescence intensity of Olig2 and PDGFRα, slices scanned with the AxioScan Z1 were analyzed using ZEN 3.1 (blue edition, Zeiss) software. When the Olig2 fluorescence intensity was comparable to background value (between 1x10^2^ and 1x10^3^), it was considered as Olig2^neg^ OPC (Supplementary Fig. 1A). To account for variability in the staining from different animal, all values of Olig2 and PDGFRα were directly compared with values of the background in the same brain section.

### Single cell RNA sequencing analysis

Source data from (Marques et al., 2016) has been used for analysis. Read count data and meta data were downloaded from the USCS Cell Browser (https://mouse-oligo-het.cells.ucsc.edu). Counts were filtered and normalized using Scanpy (Wolf, Angerer, & Theis, 2018). Cells were identified as OPCs based on the “OPC” annotation in the meta data with high *Pdgfra* expression and no *Plp1* expression. Differential expression was identified by the *scanpy.tl.rank_genes_groups* command and genes with P value <0.01 were subjected to DAVID online tool (d. W. Huang, Sherman, & Lempicki, 2009a, 2009b) to search for enriched GO terms. For Olig2^neg^ cells, only 7 genes (P<0.01) were selected which were not adequate for GO analysis (Supplementary File 1).

### Statistical analysis

Data were analyzed with Graphpad Prism 9.0 and figures were generated with Adobe Indesign 2022. At least three animals were analyzed per group and the statistical parameters are indicated in the figure legends. Data were shown as mean ± SEM.

## Acknowledgement

The authors are grateful to Daniel Schauenburg and Frank Rhode for their excellent animal husbandry and technical assistance. We thank Prof. Dr. Mengsheng Qiu (Institute of life sciences, Hangzhou Normal University, Hangzhou, China) for providing feedback on the manuscript. We are grateful to Dr. Hongkui Zeng (Allen Institute for Brain Science, Seattle, Washington, USA) for providing Rosa26-tdTomato reporter mice, Dr. Amit Agarwal (Institute for Anatomy and Cell Biology, Heidelberg University, Heidelberg, Germany) for providing Rosa26-GCaMP3 mice and Prof. Dr. Jacqueline Trotter (Molecular Cell Biology, Johannes-Gutenberg University, Mainz, Germany) for providing AN2/NG2 antibodies.

## Supplementary Figure legends

**Supplementary Figure 1.** Observation of PDGFRα^pos^Olig2^neg^ cells in the p14 mouse brain. **A**, Pα^pos^ cells can be classified in two subgroups according to their Olig2 immunoreactivity. **B**, Immunostaining of Pα and Olig2 in the CA1 region of p14 mouse brain. **C**, Quantification of Pα^pos^Olig2^neg^ and Pα^pos^Olig2^pos^ cell density and proportion of Olig2^neg^ cells in the subregions of CA1 region, including stratum oriens (so), stratum pyramidale (sp) and stratum radiatum (sr) (indicated as blue, green and magenta in **D**, respectively).

**Supplementary Figure 2.** Pα^pos^Olig2^neg^ cells are *bona fide* OPCs. **A-G**, Pα^pos^Olig2^neg^ cells of CC, CA1 and DG were co-stained with markers of OPCs (NG2, **A**), oligodendrocyte lineage cells (Sox10, **B**), mature oligodendrocytes (CC1, **C**), astrocytes (GFAP, **D, E**), microglia (Iba1, **E**), neurons (NeuN, **F**), or precursor markers (Sox2, **H**). Arrowheads: Olig2^pos^ OPCs, arrows: Olig2^neg^ OPCs.

**Supplementary Figure 3.** Pα^pos^Olig2^neg^ cells are neither apoptotic cells nor phagocytosed by microglia in CC, CA1 and DG. **A**, Pα^pos^Olig2^neg^ cells did not express apoptotic markers. **B**, Pα^pos^Olig2^neg^ cells were not phagocytosed by microglia. Arrowheads: Olig2^pos^ OPCs, arrows: Olig2^neg^ OPCs.

**Supplementary Figure 4.** Pα^pos^Olig2^neg^ cells do not express Ki67, while frequently found along the brain vasculature. Arrowheads: Olig2^pos^ OPCs, arrows: Olig2^neg^ OPCs.

**Supplementary Figure 5.** Olig2^neg^ OPCs are induced by ischemic insults in the hippocampus of mice with middle cerebral artery occlusion (MCAO). **A, B**, Immunostaining of GFAP (**A**) and Iba1 (**B**) after 3 days post MCAO indicates intensive gliosis in the hippocampus. **C**, Magnified views of the hippocampus stained with Pα and Olig2. **D**, Cell density of Olig2^neg^ OPCs were increased at the ipsilateral side compared to the contralateral side. Arrowheads: Olig2^pos^ OPCs, arrows: Olig2^neg^ OPCs.

**Supplementary Figure 6.** Complex motor learning does not change the density of Pα^pos^Olig2^neg^ cells in the motor cortex or corpus callosum. **A**, Experimental schedule for testing motor performance employing the Erasmus Ladder. Mice at the age of 8 weeks or 19 weeks were trained running on the Erasmus Ladder for three weeks and analyzed at before, during or after the learning. **B,** Percentage of missteps decreased after two weeks of training indicating successful learning. **C**, Immunostaining of PDGFRα (Pα) and Olig2 in hippocampus before or during the learning. **D, E,** Quantification of Pα^+^Olig2^neg^ cells in DG and CA1 of control (ctl), before, during and after learning sessions. **F, G**, Immunostaining of Pα and Olig2 in the motor cortex (mCTX) and corpus callosum (CC) of control (**F**) and mice after learning (**G**). **H-J**, Comparison of the Olig2^neg^ OPCs density in somatosensory CTX (**H**), CC (**I**) and mCTX (**J**) with or without motor learning at the age of 8 weeks or 19 weeks. Arrowheads: Olig2^pos^ OPCs, arrows: Olig2^neg^ OPCs.

---

**Supplementary table 1.** Primary antibodies.

| Antibodies | Host | Dilutions | Cat. No. | Company | PRID |
| --- | --- | --- | --- | --- | --- |
| PDGFRα | goat | 1:500 | AF1062 | R&D Systems | AB_2236897 |
| adenomatous polyposis coli clone 1 (CC1) | mouse | 1:200 | OP80 | Calbiochem | AB_2057371 |
| Olig2 | rabbit | 1:500 | AB9610 | Millipore | AB_570666 |
| Olig2 | mouse | 1:500 | MABN50 | Millipore | AB_10120130 |
| NG2 | rat | 1:50 |  | Trotter Lab | AB_10807410 |
| Neun | mouse | 1:500 | MAB377 | Millipore | AB_2298772 |
| Doublecortin | G P | 1:500 | AB2253 | Millipore | AB_1586992 |
| Sox10 | rabbit | 1:500 | ab27655 | abcam | AB_778021 |
| GFAP | rabbit | 1:1000 | Z 0334 | Dako | AB_10013382 |
| Sox2 | mouse | 1:200 | MAB2018 | R&D systems | AB_358009 |
| Ki67 | rat | 1:500 | 14-5698-82 | Invitrogen | AB_10854564 |
| Cleaved Caspase-3 | rabbit | 1:200 | 9661 | Cell Signaling Technology | AB_2341188 |
| BrdU | rat | 1:100 | 2313786 | abcam | AB_2341188 |
| CD31 | rat | 1:100 | MAB5406 | BD Pharmingen | AB_393571 |
| GFP | chicken | 1:1000 | A10262 | Invitrogen | AB_2534023 |
| Iba1 | rabbit | 1:1000 | 019-19741 | Wako | AB_839504 |
| CD68 | rat | 1:500, | MCA1957GA | Bio Rad | AB_324217 |

**Supplementary table 2.** Secondary antibodies.

| **Antibodies** | **Fluorophore** | **Cat. No.** | **Company** |
| --- | --- | --- | --- |
| Donkey anti-mouse | Alexa Fluor® 488 | A21202 | Thermo Fisher |
|  | Alexa Fluor® 546 | A10036 |  |
|  | Alexa Fluor® 647 | A31571 |  |
|  | DyLight® 755 | SA5-10171 | Invitrogen |
| Donkey anti-rabbit | Alexa Fluor® 488 | A21206 | Thermo Fisher |
|  | Alexa Fluor® 546 | A10040 |  |
|  | Alexa Fluor® 647 | A31573 |  |
|  | Alexa Fluor® 790 | A11374 |  |
| Donkey anti-goat | Alexa Fluor® 488 | A11055 | Thermo Fisher |
|  | Alexa Fluor® 546 | A11056 |  |
|  | Alexa Fluor® 647 | A21447 |  |
|  | Alexa Fluor® 750 | ab175744 | abcam |
| Donkey anti-rat | DyLight-755 | SA5-10031 | Thermo Fisher |
| General stain for nuclei | DAPI (25 ng/ml) | A10010010 | BioChemica |

