## Supplementary figures and images for "Brain injuries and complex motor learning suppress Olig2 in a subpopulation of oligodendrocyte precursor cells"

### supplementary figures

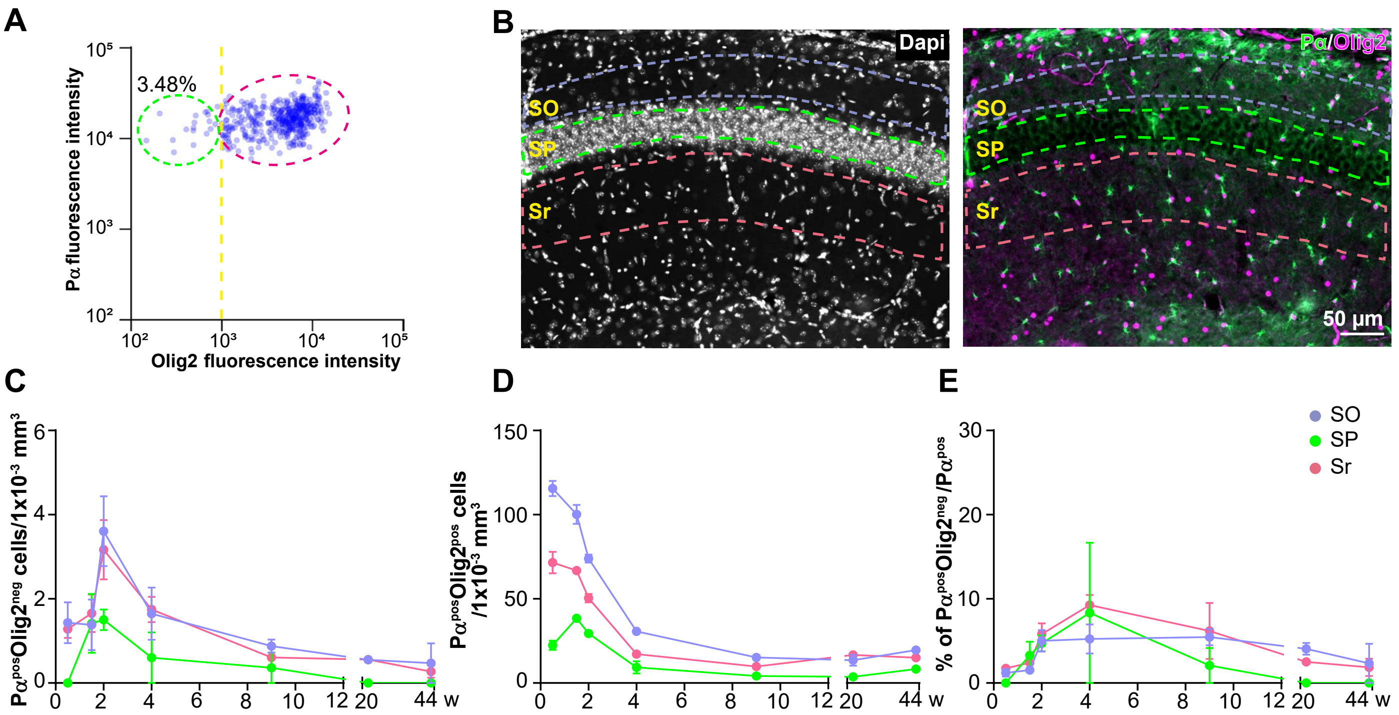

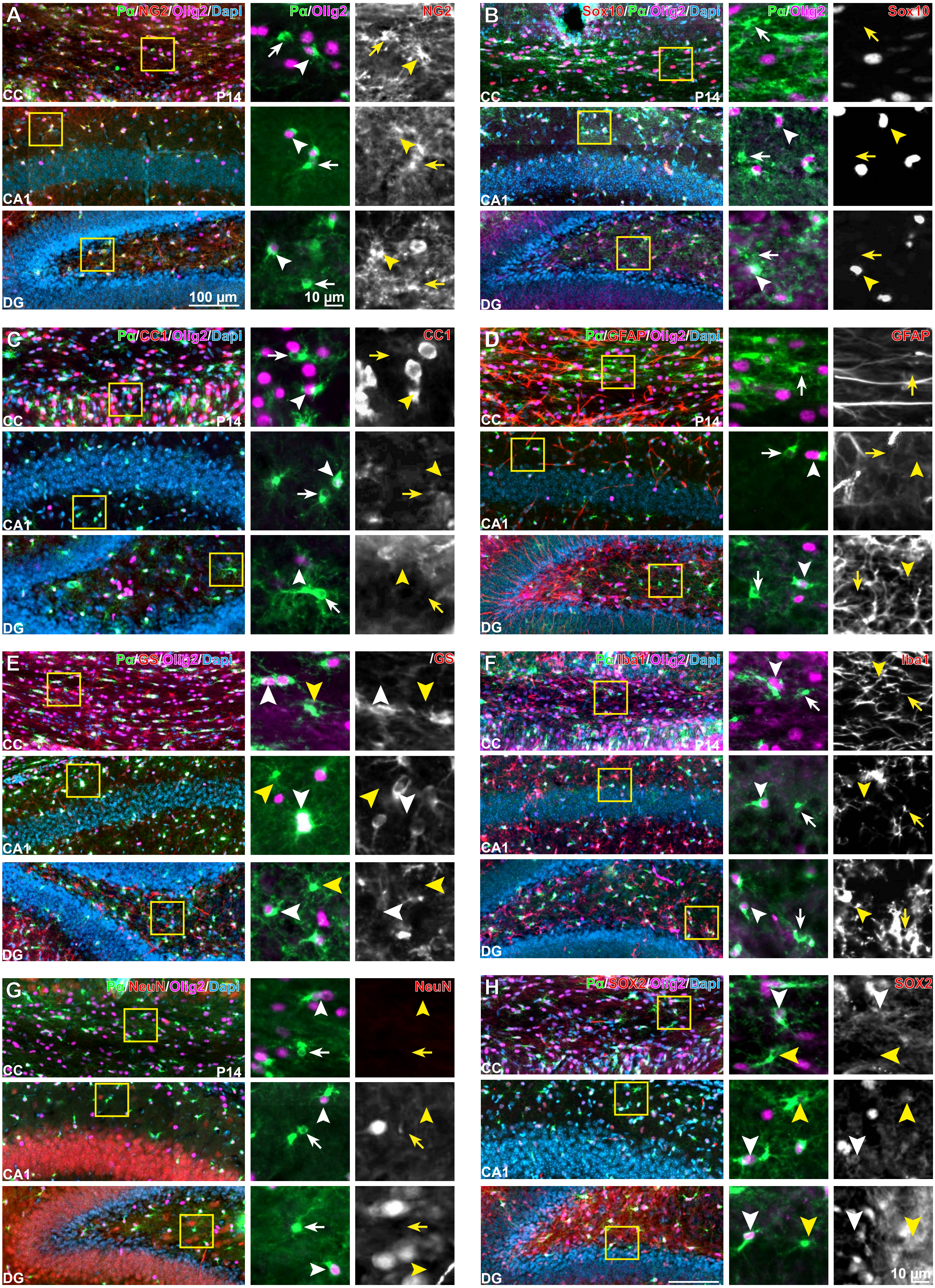

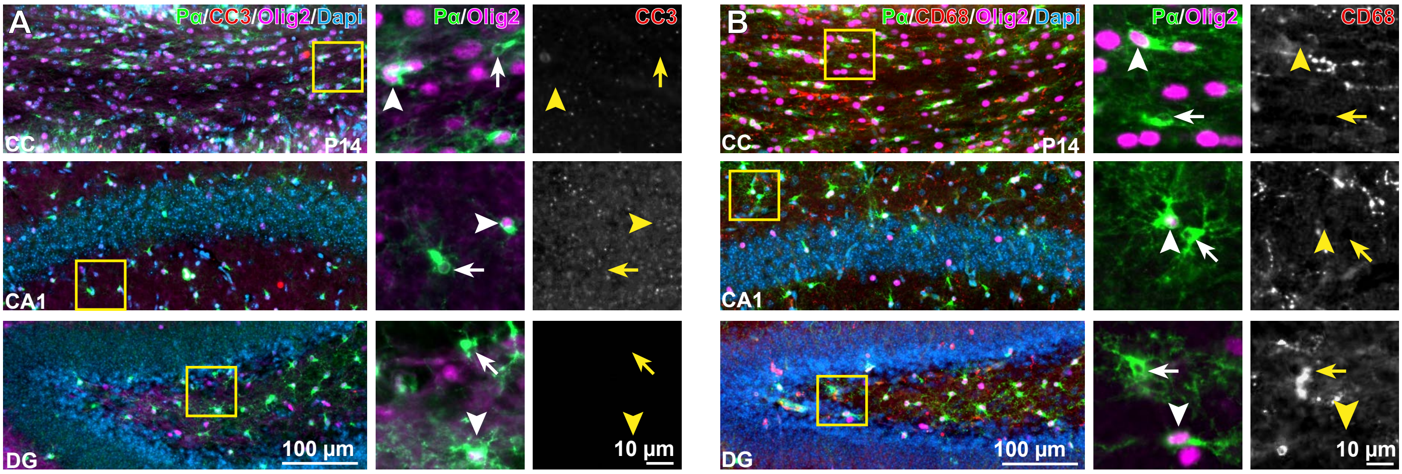

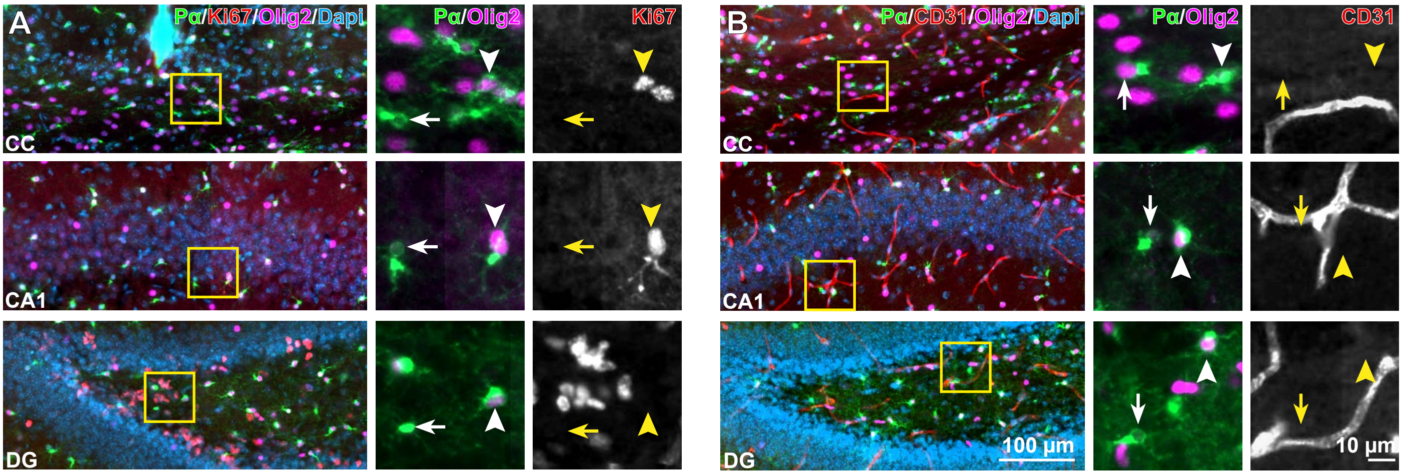

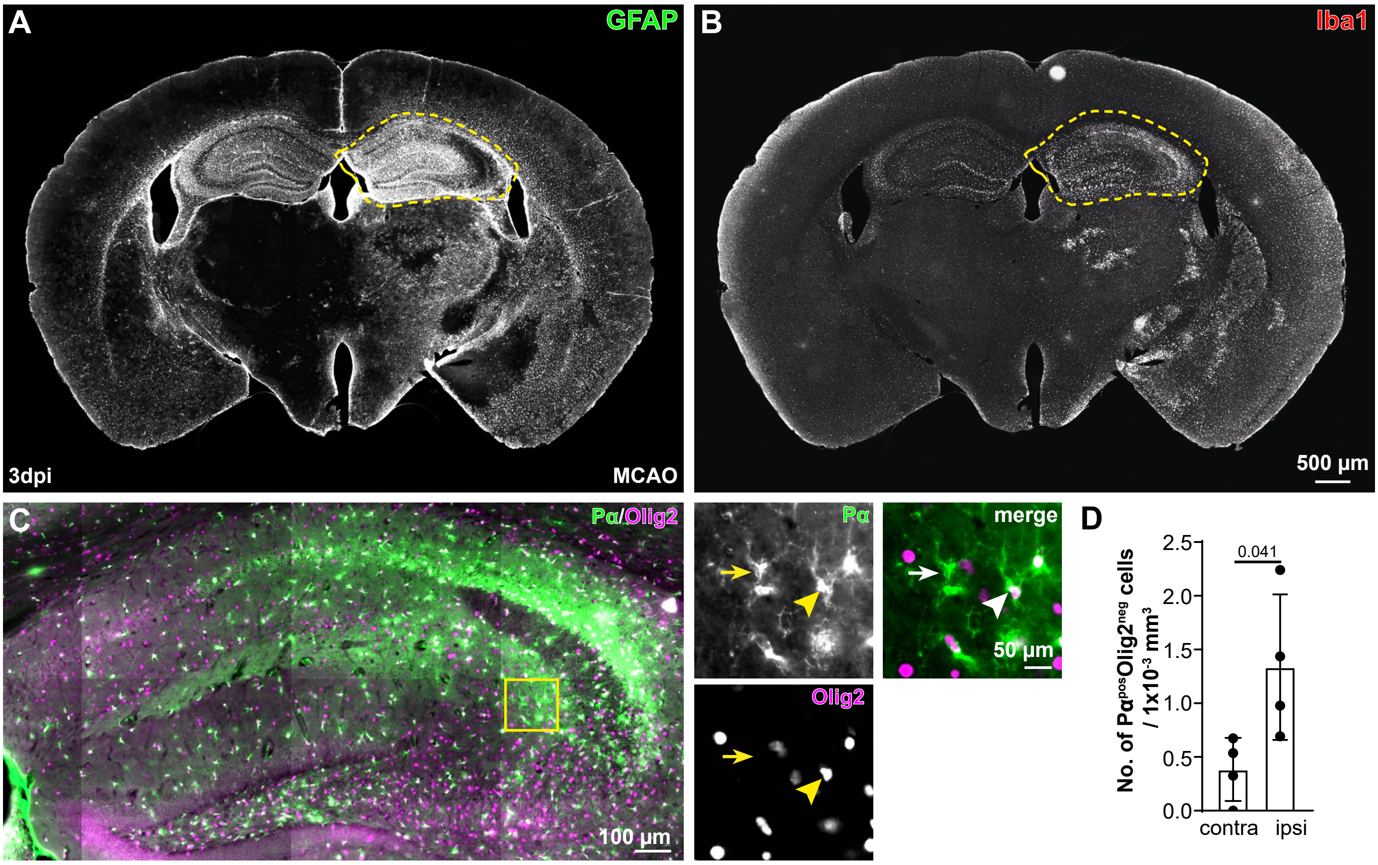

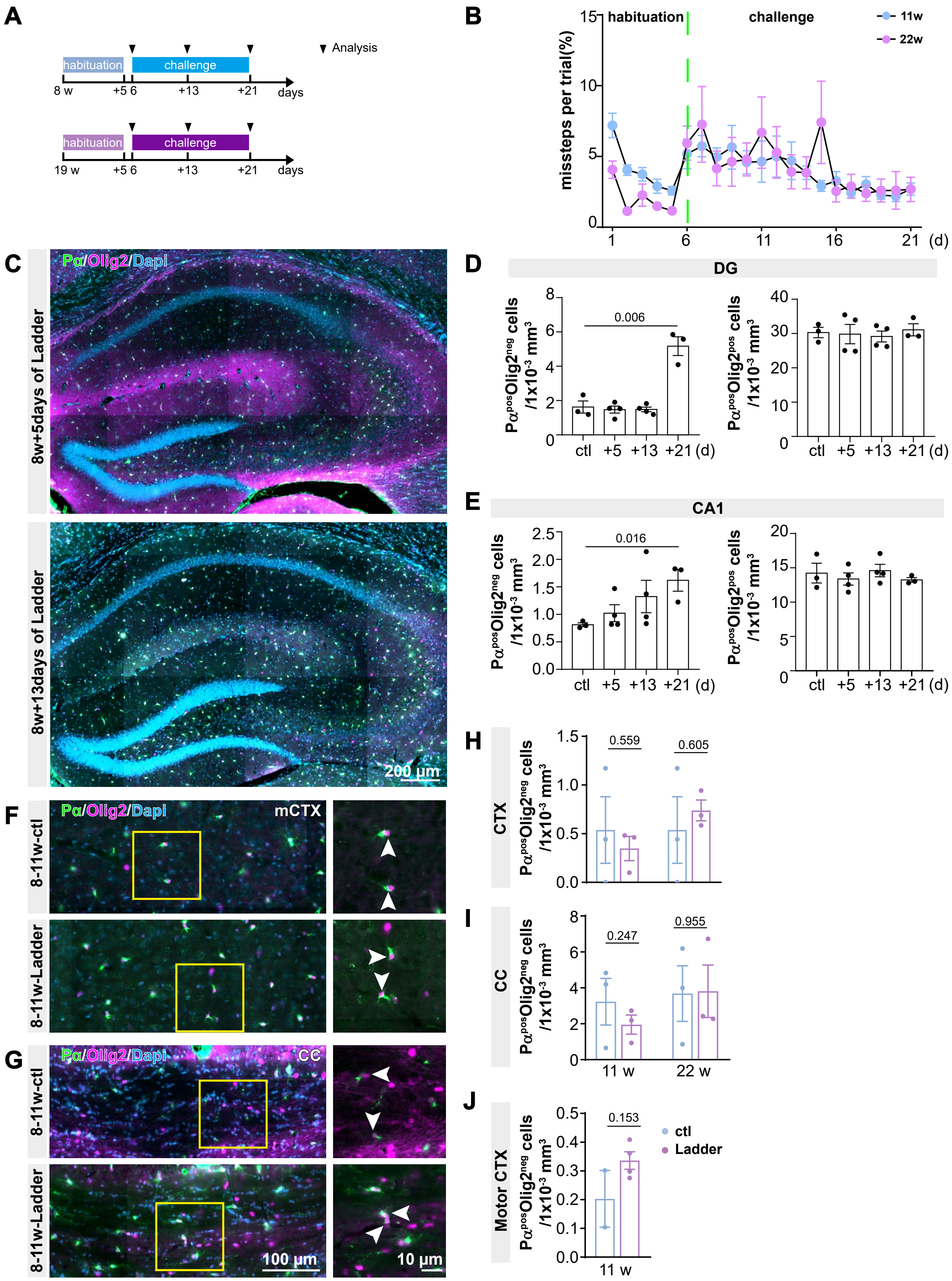
